# Cyclin C nuclear release and mitochondrial dysfunction define molecular signatures of *MED13L* Syndrome

**DOI:** 10.64898/2026.06.01.729270

**Authors:** Alicia N. Campbell, Kendall N. Jung, Steven J. Doyle, Ekaterina Lebayle, Barbara Corneo, Joanna Feng, Christopher L. Ricupero, Jennifer M. Bain, Randy Strich

## Abstract

The Mediator Kinase Module (MKM) coordinates transcriptional programs regulating cellular metabolism, stress responses, and differentiation. Heterozygous variants of *MED13L*, a core MKM component, cause a neurodevelopmental disorder characterized by variable intellectual disability, developmental delay, neuromuscular dysfunction, and congenital anomalies. However, the molecular basis underlying this clinical heterogeneity is poorly defined. Previously, we identified mitochondrial dysfunction and aberrant nuclear release of another MKM component, cyclin C (CCNC), in a single *MED13L* syndrome patient-derived fibroblast line. Here, we expand these studies across 12 patient-derived fibroblasts harboring 11 distinct *MED13L* variants. We identify mitochondrial dysfunction as a consistent feature of *MED13L* variation, characterized by reduced mitochondrial ATP production, decreased mitochondrial DNA abundance, elevated reactive oxygen species, and impaired transcription of genes involved in mitochondrial biogenesis. In parallel, all variant lines exhibit aberrant cytoplasmic CCNC localization, consistent with its established role in mitochondrial fission. Longitudinal analyses further reveal progressive declines in mitochondrial function associated with premature cellular aging, consistent with increased metabolic deficits. Importantly, the severity of mitochondrial dysfunction shows an association with variant position within *MED13L* and with clinical functional measures, suggesting that mutation location may partially predict disease severity. Together, these findings establish mitochondrial dysfunction as a consistent cellular feature of *MED13L* heterozygosity and identify CCNC mis-localization as a candidate biomarker of MKM disruption. More broadly, this work reveals an intersection between transcriptional control and mitochondrial homeostasis in *MED13L* syndrome, forming the framework for biomarker-driven therapeutic development in *MED13L*-associated and related neurodevelopmental disorders.

## Introduction

*MED13L* Syndrome is a rare neurodevelopmental disorder characterized by global developmental delay, hypotonia, congenital heart defects, seizures, and variable intellectual disability^1–4^ (*MED13L* Syndrome MIM #616789; *MED13L* MIM #608771). Despite increasing identification of pathogenic variants in *MED13L* through clinical sequencing, substantial clinical heterogeneity remains, even among individuals with similar mutations^4,5^. This variability has complicated efforts to define disease mechanisms and has limited the development of targeted therapies.

*MED13L* encodes a core structural component of the Mediator Kinase Module (MKM), which also includes its paralog MED13, CDK8 or CDK19, cyclin C (CCNC), and MED12 or MED12L^6–8^. The MKM acts as a regulatory center that integrates transcriptional control with cellular stress responses and differentiation cues, enabling rapid transcriptional reprogramming during developmental transitions and metabolic adaptation^8–12^. Beyond its canonical nuclear role, emerging evidence suggests that MKM activity is linked to mitochondrial regulation through stimulus-dependent re-localization of CCNC, which has been implicated in mitochondrial fission and cellular stress responses^13–16^. Under mitochondrial or oxidative stress, a portion of CCNC translocates from the nucleus to the cytosol, where it promotes mitochondrial fission, thereby coupling transcriptional reprogramming to organelle quality control^14,17,18^. Disruption of this coordinated signaling axis has the potential to impair both nuclear transcriptional programs and mitochondrial homeostasis.

Our previous study of *MED13L* disruption identified transcriptional dysregulation and mitochondrial abnormalities in limited cellular models^15^. Previous RNA-seq analyses from *Ccnc^-/-^*mouse embryonic fibroblasts (MEFs) identified a transcriptional network governing mitochondrial maintenance, including genes involved in mitochondrial biogenesis, transcription, and nuclease activity^19,20^. However, these findings have been derived from small cohorts, and it remains unclear whether mitochondrial dysfunction represents a consistent and defining feature of *MED13L* syndrome or reflects variant-specific effects.

Importantly, while clinical reports frequently describe metabolic abnormalities in individuals with *MED13L* syndrome, including elevated lactate, reduced exercise tolerance, and features suggestive of altered bioenergetic regulation, systematic cellular phenotyping across diverse genotypes has not been performed^5,21–24^. Establishing whether mitochondrial dysfunction represents a consistent and distinguishing cellular feature of *MED13L* dysfunction would provide critical mechanistic insight and could enable the development of cellular biomarkers for diagnosis and treatment monitoring. Moreover, whether such defects represent static abnormalities or progressive vulnerability has not been examined.

Here, we examine a panel of patient-derived fibroblast lines carrying heterozygous deletion, nonsense, frameshift, and missense variants in *MED13L* to define shared cellular features. We identify mitochondrial dysfunction as a reproducible phenotype characterized by impaired ATP production, reduced mitochondrial DNA content, elevated oxidative stress, and altered transcriptional programs governing mitochondrial maintenance. In parallel, we observe aberrant cytoplasmic localization of CCNC across variant cell lines, suggesting disruption of MKM-associated signaling. In addition, we observed premature acquisition of senescence-associated phenotypes. Together, these findings implicate mitochondrial dysfunction as a previously underappreciated cellular feature of *MED13L* variation and nominate CCNC mis-localization as a candidate cellular biomarker of MKM disruption and mitochondrial stress. This work establishes a functional link between Mediator-associated transcriptional regulation and mitochondrial dynamics, thus expanding the conceptual framework for *MED13L* syndrome pathophysiology.

## Materials and Methods

### Human subjects and fibroblast biopsies

Consent was granted according to IRB protocols (IRB #AAR7203, Columbia University Irving Medical Center). Individuals ranged in age from 17 months to 17 years and included six males and six females. At the time of collection, clinical severity metrics, including Gross Motor Function Classification System (GMFCS) communication scores, were collected and scores ranged from 1-5 where available^25^. Control fibroblasts were purchased from ATCC (ATCC: Primary Dermal Fibroblasts Normal; Human, Neonatal (HDFn): PCS-201-010). All patient-derived lines carried heterozygous pathogenic or likely pathogenic variants in *MED13L,* and included missense, frameshift, nonsense, and exon-deletion alleles annotated from clinical genomic analysis. Variant classification (IDR, N-terminal, C-terminal and medPIWI) were ascribed based on UniProt functional domains (MED13L_Human #Q71F56). *MED13L* syndrome (MIM #616789; ICD-10-CM Q87.85) was diagnosed in individuals with a pathogenic or likely pathogenic heterozygous mutation in the *MED13L* gene (MIM #608771), located on chromosome 12q24.21. Primary dermal fibroblasts were obtained from individuals diagnosed with *MED13L* syndrome through institutional biobanking protocols at Columbia University Irving Medical Center, Stem Cell Initiative. Detailed protocols can be found in Iannello et al., 2023 (PMID: 36912580)^26^. Briefly, fibroblast medium was prepared containing DMEM high glucose (Life Technologies, 11960077), 10% heat-inactivated fetal bovine serum (FBS) (Life Technologies, 10082147), 1X penicillin-streptomycin (Life Technologies, 15140122), 1X L-glutamine (Life Technologies, 25030081), and 1X sodium pyruvate (Life Technologies, 11360070), and sterile filtered. Biopsies were collected on either the inner upper arm or lateral upper thigh. A ∼3-mm biopsy was collected and placed in prewarmed fibroblast medium.

One well of a 6-well tissue culture dish was treated with 0.1% gelatin (Millipore Sigma, SF008) for 30m at 37°, 5% CO_2_. Gelatin coating was aspirated, and biopsy was added to the well. Biopsy was pinned to the culture dish using 30-G needle. Warmed fibroblast medium was added to the well, and lid replaced. The plate was transferred to standard cell culture conditions with fibroblast medium replaced every 3-4 days.

### Culturing dermal fibroblasts

Culturing of initial fibroblasts was completed by CUIMC Stem Cell Initiative and based on a published protocol^26^. Briefly, once the initial fibroblast layer was fully confluent, the medium was aspirated and additional culturing dishes were prepared with gelatin as before. Cells were washed with 1X DPBS (Corning, MT-21-030-CV). Cells were then released from the dish using 0.25% trypsin-EDTA (Life Technologies, 25200072) and incubated for 2 min at 37°. Trypsin was quenched with fibroblast medium (see protocol above) and resuspended using pipette. Cells were collected in 15 mL tubes, and an additional 10 mL medium was added before centrifugation for 5 min at 300 x *g*, RT. The supernatant was aspirated, and the cell pellet was resuspended in 1 mL medium, followed by an additional 11 mL. The cell suspension was dispersed across 12-well plates and cultured under conditions previously described.

### Characterization of dermal fibroblasts grown from skin biopsy

The fibroblasts were routinely tested for the absence of mycoplasma contamination using a PCR-based kit (e-Myco plus, Intron, 25234). For cell characterization and staining, cells were washed with 1X DPBS (Corning, MT-21-030-CV), and fixed with 4% PFA (Santa Cruz, sc-281692) for 10 min, RT. Cells were washed once more, then permeabilized with 0.1% Triton X-100 (Thermo Fisher Scientific, 85111) in 1X PBS. Cells were blocked with protein blocking buffer (Agilent, X0909) for 30 min at RT. Cells were stained with appropriate primary and secondary antibodies (see previous publication for full antibody list, dilutions, and manufacturers). Cells were analyzed using fluorescence microscopy.

### General Cell Culture

Fibroblasts were maintained in Dulbecco’s modified Eagle’s medium (DMEM) (Corning, MT10013CV) supplemented with 10% FBS (Avantor, MP1300500) and 1% penicillin/streptomycin (HyClone, 16777-164) at 37°C and 5% CO_2_. Media was replenished every 48-72 hr, and cells were passaged using 0.25% Trypsin-EDTA (Gibco, 25-200-114) when reaching approximately 80-90% confluency.

### Variant annotation and classification

Transcript annotations were based on Ensembl GRCh38 reference sequences. Frameshift variants were annotated by incorporating the number of altered nucleotides and predicted downstream translation distance using NMDetective conventions^27^. Missense variants were independently classified using MutPred-LOF^28^. Exon-level deletion variants were classified based on predicted effects on transcript structure and translation. Deletions involving one or two consecutive exons were designated small exon deletions; those spanning three or more exons were designated large exon deletions. Deletion variants were evaluated for predicted transcript stability and proximity to the terminal exon junction rather than the predicted reading frame alone. The large exon 3-25 deletion removes the majority of the coding region and was therefore considered a severe truncating allele, and its precise transcript fate could not be confidently determined.

### RNA Isolation and quantitative PCR

Total RNA was extracted using column-based purification with on-column DNase digestion (NEB, 103529-148). cDNA was synthesized using kit (ThermoFisher, FERK1642). qPCR was performed using SYBR green (Applied Biosystems, 4368702) and machine (Applied Biosystems, 43-766-00). Gene expression was normalized to the housekeeping gene *GAPDH* and calculated using the ΔΔC_t_ method^29^. Transcript abundance was expressed as log_2_ fold change relative to pooled control fibroblasts. Target gene primers can be found in Table S2.

### Immunofluorescence Microscopy

Cells were seeded on glass coverslips and grown overnight. The following day, cells were stained with MitoTrackerRed (Invitrogen, M7512) and fixed with 4% paraformaldehyde. Permeabilization was performed using 0.2% Triton X-100, followed by blocking in serum-based buffer (Licor, NC1660556). Primary antibodies were added and incubated overnight at 4°C. Antibodies included dsDNA (DSHB, AB10805293) and CCNC (CST, 68179). Fluorophore-conjugated secondary antibody goat anti-rabbit IgG or goat anti-mouse IgG was applied the next day following washing (Invitrogen, A11008; Invitrogen, A32723). Nuclei were labeled with mountant containing DAPI. Images were acquired using fluorescence microscopy at identical exposure settings for all conditions using a Keyence BZ-X 710 Inverted Fluorescence Microscope.

### Quantification of Mitochondrial Fragmentation

Mitochondrial morphology was scored blind to genotype and assessed by three separate researchers. Cells were classified as “fragmented” if >50% of the mitochondrial network consisted of punctate or short tubular structures rather than elongated networks. At least 100 cells per cell line were analyzed across replicate experiments, and the percentage of cells exhibiting fragmentation was calculated. See Figure S1 for examples of mitochondrial morphology that were used for quantification purposes.

### Seahorse mitochondrial and glycolytic ATP production rate assays

Cellular bioenergetics were quantified using the Seahorse XFe24 or XFp Real-Time ATP Rate Assay. Fibroblasts were seeded into plates at a density of 20,000 cells per well and incubated overnight in standard conditions. On the day of the assay, cells were washed and incubated in Seahorse XF assay medium supplemented with glucose, glutamine, and sodium pyruvate and equilibrated in a non-CO_2_ incubator for 45-60 min prior to analysis. Basal oxygen consumption rate (OCR) and proton efflux rate (PER) values were recorded prior to compound injection. Sequential injections were performed according to the manufacturer’s protocol.

### Seahorse Normalization and Quality Control

ATP production values were normalized to cell number. Wells containing abnormal OCR/PER traces, loss of adherence, or incomplete drug response were excluded based on Seahorse quality control flags. Each condition/cell line was run in technical duplicates with a minimum of three biological replicates.

### Calculation of mitochondrial and glycolytic ATP production rates

The OCR and extracellular acidification rate (ECAR) values were exported from Wave software and converted to ATP production rates according to Agilent Seahorse Real-Time ATP Rate Assay calculations with plate-specific correction factors. Briefly, ECAR values were first converted to PER by averaging the first three baseline ECAR measurements and applying assay-specific buffer and well volume correction factors. Conversion constants were applied as instructed by the manufacturer and adjusted based on the instrument used (XFp or XFe24). Basal mitochondrial oxygen consumption (OCR_basal_) was calculated as the average of the first three baseline OCR measurements. Non-mitochondrial respiration was defined as OCR remaining after rotenone/antimycin A injection and was subtracted from OCR_basal_ to yield mitochondrial-derived oxygen consumption (mitoOCR). To account for mitochondrial-driven extracellular acidification, mitochondrial PER (mitoPER) was calculated as 0.5x mitoOCR, and this value was subtracted from total PER to obtain glycolytic PER (glycoPER). Glycolytic ATP production (glycoATP) was calculated directly from glycoPER after correction using assay-specific conversion factors. Mitochondrial ATP production (mitoATP) was calculated from the oligomycin-sensitive fraction of respiration. OCR following oligomycin addition was averaged across three timepoints and subtracted from OCR_basal_ to determine ATP-linked respiration (OCR_ATP_). OCR_ATP_ values were converted to ATP production rates using manufacturer-provided P/O ratios and plate-specific assay constants. Total ATP production was calculated as the sum of mitochondrial and glycolytic ATP production rates. All ATP values were reported as pmol ATP/min and normalized to cell number.

### Statistical analysis of longitudinal Seahorse Data

To evaluate whether mitochondrial ATP production declined progressively with serial passage, time-course Seahorse data were analyzed using nonlinear regression with a straight-line model in GraphPad Prism. For each cell line, mitochondrial ATP production values across generations were fit using least squares regression. Global curve comparison was performed using extra sum-of-squares F-tests to determine whether slopes or Y-intercepts differed between control and *MED13L* variant cells. Separate comparisons were conducted for slope (rate of change over time) and y-intercept (baseline mitoATP). Shared-parameter models were compared against independent fits to assess whether slopes or intercepts were significantly different between datasets.

### Senescence-associated β-galactosidase assay

Senescence was evaluated using a fluorogenic β-galactosidase reporter assay (Invitrogen, C10851). Cells were stained according to manufacturer’s instructions and analyzed via BD Accuri C6 Plus flow cytometer. Values were analyzed and background-subtracted fluorescence intensity was calculated for each sample.

### BrdU Incorporation assay

Cellular proliferation was quantified using bromodeoxyuridine (BrdU) incorporation followed by fluorogenic detection using flow cytometry. Patient-derived *MED13L* fibroblasts and control fibroblasts were seeded at equivalent densities and grown under standard culture conditions until ∼60-70% confluency. Cells were pulsed with BrdU (final concentration 10 μM) for 24 hr to label cells actively undergoing DNA synthesis. Following labeling, cells were washed with PBS then fixed and permeabilized using 4% paraformaldehyde and 0.2% Triton X-100, respectively. DNA was chemically denatured using 2N HCl, and BrdU detected using BrdU antibody (Invitrogen, B35130) with overnight incubation at 4°C. BrdU signals were acquired on BD Accuri C6 Plus flow cytometer as described above. All experiments were performed using at least three independent biological replicates per genotype with technical duplicates.

### Cytosolic reactive oxygen species (ROS) measurement

Cytosolic ROS was measured using dihydroethidium (DHE), which fluoresces upon oxidation by superoxide. Fibroblasts were seeded into 6-well plates and harvested on the day of the assay according to normal detachment protocol (see above). Cell suspensions were incubated with DHE at 37°C for 20-30 min protected from light. Cells were then washed and analyzed via flow cytometry. Single-cell suspensions were gated based on forward and side scatter to exclude debris and doublets. Median fluorescence intensity was calculated for each population and normalized to matched control samples included on each run, and background fluorescence determined by unstained controls.

### Mitochondrial ROS measurement

Mitochondrial superoxide production was quantified using MitoSOX Red. Cells were grown in 6-well plates, harvested, and incubated with MitoSOX at 37°C for 30 min protected from light. Cells were washed and analyzed immediately via flow cytometry as described above. All ROS measurements were obtained from a minimum of three independent experiments with several technical replicates within each run.

### Mitochondrial DNA copy number

Total DNA was extracted using Omega Bio-Tek EZNA Tissue DNA Kit (NC0721312). mtDNA copy number was quantified using qPCR with primers targeting mitochondrial gene loci and nuclear reference genes (see Table S2 for primers). Each reaction was run in triplicate with a minimum of two technical duplicates. mtDNA copy number was calculated by normalizing mitochondrial targets to nuclear DNA and expressing values relative to controls.

### Clinical and behavioral data collection

Caregiver-reported clinical and behavioral data were obtained from two rare disease research registries: Simons Searchlight and RARE-X. Simons Searchlight is an international natural history and genetics research program that collect standardized phenotypic, behavioral, and developmental information from individuals with rare neurodevelopmental disorders and their caregivers, and is sponsored by the Simons Foundation Autism Research Initiative. RARE-X is a patient-driven data collection platform that combines longitudinal clinical and caregiver-reported outcomes across rare disease populations^30^. Data from both registries were harmonized to generate unified clinical variables for downstream analyses.

### Composite Functional/Behavioral Score

The Composite Functional/Behavioral Score, utilized adaptive functioning metrics derived from the Vineland Adaptive Behavior Scales, Third Edition (Vineland-3; VABS-3)^31^ when available from either registry. Three behavioral domains were incorporated into the composite metric: Adaptive Behavior Composite (ABC), communication, and motor functioning scores. From Simons Searchlight, the Vineland-3 Adaptive Behavior Composite standard score was obtained directly from the ABC standard score field. From RARE-X, the corresponding value was obtained from the vi3_adaptive_composite variable within the Vineland dataset. ABC scores were additionally categorized into severity bins based on current clinical standards: mild (>85), borderline (70-84), or moderate/severe (<70).

To generate the composite metric encompassing all three, each domain score was normalized by z-score standardization using the cohort mean and standard deviation. Typically, higher ABC scores reflect improved adaptive functioning, whereas the remaining clinical measures were oriented such that increasing values reflected worsening impairment. Therefore, normalized ABC score was directionally inverted prior to composite score generation to maintain consistency across variables. Communication domain scores were obtained from the Vineland-3 communication domain in either registry and similarly normalized. Motor scores were derived from Vineland-3 gross and fine motor assessments (broad “motor” category) and normalized in the same manner.

The final composite functional score was calculated by summing all available normalized domain scores for each individual and dividing by the number of reported domains to account for incomplete datasets or missing variables. Severity thresholds were selected based on conventional interpretation ranges for standardized Vineland-3 domain scores, which are normed to a mean of 100 and standard deviation of 15^32^. This approach enabled incorporation of participants with partial phenotypic data while preserving relative weighting across behavioral domains.

### Autism and Social Trait Characterization

Autism and social impairment traits were harmonized across caregiver-reported datasets as described above. Here, datasets were combined to generate an ordinal Autism/Social Trait severity score ranging from 0-3. Variables were selected based on overlaps in social functioning and autism-related behavioral constructs between registries.

Autism spectrum disorder (ASD) diagnosis status was first determined using either registry reported clinical diagnoses or validated autism-related behavioral assessments. Within Simons Searchlight, ASD status was derived from Social Communication Questionnaire (SCQ) and/or Social Responsiveness Scale (SRS) clinical cutoffs or reported diagnoses. Within RARE-X, ASD diagnosis status was obtained from the medical history/diagnosis dataset (ASD_Diagnosis). When available, clinician-reported ASD diagnoses were prioritized over questionnaire-derived classifications. ASD diagnosis was recorded as a binary variable (yes/no).

To harmonize measurements across registries, SCQ scores were categorized into ordinal severity tiers with predefined cutoff ranges: scores <10 were assigned Tier 0, scores 10-14 were assigned Tier 1, scores 15-21 were assigned Tier 2, and scores >21 were assigned Tier 3. SCQ scores were stratified into ordinal categories centered around established ASD screening thresholds to facilitate cross-registry analyses^33^. SRS T-scores were similarly categorized, with scores <60 assigned Tier 0, scores 60-65 assigned Tier 1, scores 66-75 assigned Tier 2, and scores >75 assigned Tier 3. Increasing tier values reflected greater autism-associated symptom severity and social impairment. SRS T-score severity tiers were assigned using established SRS-2 clinical interpretation thresholds^34^.

Within RARE-X, social impairment severity was assessed using Vineland-3 socialization domain scores (vi3_socialization). Scores were converted into ordinal severity tiers such that scores >85 were assigned Tier 0, scores 70-84 were assigned Tier 1, scores 55-69 were assigned Tier 2, and scores <55 were assigned Tier 3, with increasing severity corresponding to greater impairment in social functioning.

To align measurements across registries, SCQ severity tier, SRS severity tier, prior ASD diagnosis, and Vineland socialization severity tiers were integrated into a unified “Autism Trait Tier” ranging from 0-3. When multiple assessments were available for a single participant, clinician or caregiver-reported ASD diagnoses were prioritized over questionnaire-derived classifications. The final harmonized Autism Trait Tier was used downstream for genotype-phenotype and behavioral correlation analyses.

### Statistical analyses

All statistical analyses were performed using GraphPad Prism. All values represent biological replicates derived from independent fibroblast lines unless otherwise stated. For comparisons involving multiple variant groups relative to controls, one-way ANOVA with Dunnett’s post hoc test for multiple comparisons was applied. Linear regression was used to assess relationships between variant position and phenotype metrics. Goodness-of-fit was evaluated by using R values and slope. Time-course Seahorse data were analyzed using linear regression to assess trends across serial passage. Slopes were compared between control and variant groups. All boxplots indicate median and interquartile range. Scatter plots show individual data points with either mean or median and SEM or SD. Sample sizes were chosen based on cell availability. Investigators were blinded to genotype during the entire study, until data analysis.

## Results

### *MED13L* variants exhibit mitochondrial dysfunction

Our previous report revealed reduced mitochondrial ATP (mitoATP) production in a single *MED13L* patient fibroblast cell line (+/S1497^fs*^)^15^. To determine whether mitochondrial dysfunction was a common phenotype associated with *MED13L* syndrome patients, an additional 12 cell lines were collected and expanded. Fibroblasts were obtained from both male and female patients ranging in age from 17 months to 17 years (see Materials and Methods and Table S1 for subject details). Following expansion in culture, mitochondrial ATP production was quantified using Seahorse metabolic flux analysis. Across the cohort, mitoATP production was significantly reduced relative to the control (human foreskin fibroblasts) in all but one cell line (N1824M^fs^*), which exhibited a similar trend, but did not reach statistical significance. (Fig. 1A open box). The overall reduction in mitoATP varied, with deletion alleles (far right, shades of green) exhibiting the greatest reduction at approximately 33% of control. The remainder demonstrated ∼50% reduction in mitoATP production. These results indicate that impaired oxidative phosphorylation is consistently observed in *MED13L* syndrome cell lines, regardless of mutant class or location.

**Figure 1.**
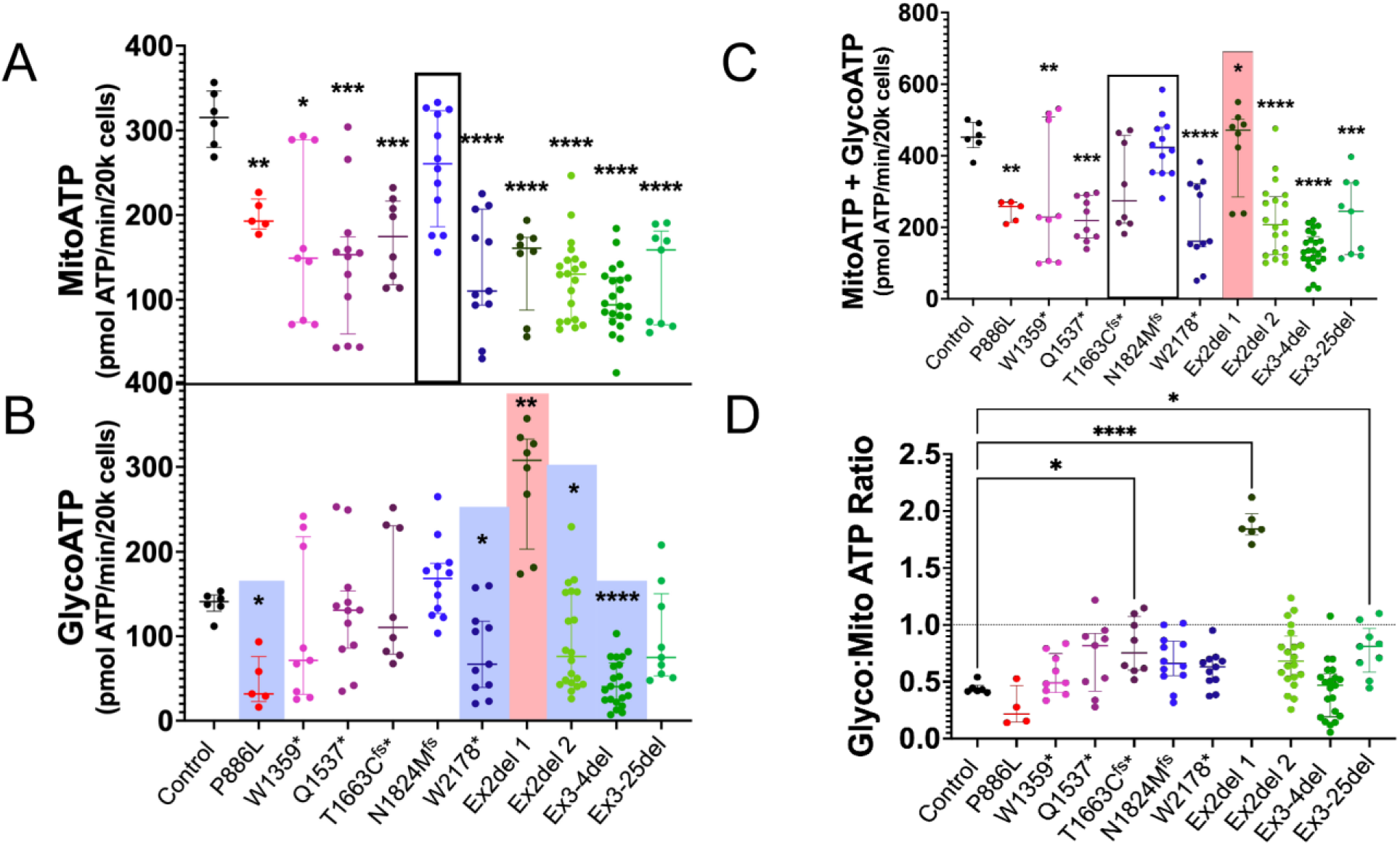
*MED13L* variants exhibit mitochondrial dysfunction. (A) Mitochondrial ATP (mitoATP) production rate as measured by Seahorse metabolic analyzer in control and *MED13L* variant fibroblasts as indicated. Open box indicates cell line with similar mitoATP production to control. (n ≥ 5 biological replicates; technical duplicates) (B) Glycolytic ATP (glycoATP) production rate measured as in (A) using proton efflux rate (n ≥ 5 biological replicates; technical duplicates). Blue boxes indicate cell lines exhibiting reduced glycoATP production, the red box indicates higher production. (C) Combined ATP production (glycoATP + mitoATP) was calculated from panels (A) and (B). Open box indicates cell lines with no difference in ATP production from control, the red box indicates higher production. The remaining cell lines exhibited reduced total ATP production. (n ≥ 5 biological replicates; technical duplicates) (D) Bioenergetic balance expressed as the ratio of mitochondrial to glycolytic ATP production (mitoATP:glycoATP) for the cell lines indicated. Values <1 indicate mitochondrial-dominant energy metabolism, whereas values approaching 1 reflect increasing reliance on glycolysis. (n ≥ 4 biological replicates; technical duplicates). In all graphs, control cell line is represented in black, missense variant in red, medPIWI/IDR mutations in shades of purple (W1359*, Q1537*, T1663C^fs*^), C-terminal variants in shades of blue (N1824M^fs^, W2178*), and exon deletions (single or multi) in shades of green (Ex2del-1, Ex2del-2, Ex3-4del, Ex3-25del). Statistical analysis was performed using one-way ANOVA with Dunnett’s post hoc correction for variant-to-control comparisons. All data represented as mean ± IQR. (*p < 0.05, **p < 0.01, ***p < 0.001, ****p < 0.0001).

Previous studies found that mitoATP production deficits are often compensated for by elevated glycolytic ATP (glycoATP) production^35–37^. Therefore, we next assessed glycoATP production in the *MED13L* variant cell lines. GlycoATP levels were variable across *MED13L* variant lines, with several exhibiting values comparable to control, while others displayed reduced glycolytic output (Fig. 1B). However, the P668L missense (red), Ex2del-2 (light green), Ex3-4del (dark green) and W2178* (dark blue) mutant cell lines exhibited reduced glycoATP production compared to control (blue boxes, Fig. 1B). Only Ex2del-1 exhibited elevated glycoATP production (red box, Fig. 1B).

To determine whether glycolytic compensation was sufficient to maintain cellular energy balance, the total ATP production was calculated as the sum of mitochondrial and glycolytic contributions (mitoATP + glycoATP). All but three cell lines exhibited significantly reduced ATP production (Fig. 1C). Two cell lines (N1824M^fs^*, T1663C^fs^*) exhibited similar ATP production to control (open box, Fig. 1C), while the Ex2del-1 cell line exhibited higher ATP production due to the elevated glycoATP production (red box). We next examined the relative contribution of glycolytic versus mitochondrial ATP production using a glyco:mito ATP ratio. While control fibroblasts derived the majority of ATP from mitochondrial sources (approximate ratio value of 0.5; indicating oxidative phosphorylation provided twice the ATP than glycolysis), several *MED13L* variant lines exhibited a shift toward increased reliance on glycolysis (Fig. 1D). However, this shift did not fully compensate for reduced mitochondrial output. Together, these findings indicate that impaired mitochondrial ATP production is a consistent feature across *MED13L* variant fibroblasts, with incomplete and variable glycolytic compensation.

### *MED13L* variants are associated with aberrant CCNC nuclear release and mitochondrial fragmentation

In unstressed cells, CCNC is predominantly localized in the nucleus, where it regulates transcription. Our previous study reported a portion of CCNC in the cytoplasm in an unstressed +/S1497^fs^* *MED13L* syndrome cell line^15^. To determine whether this phenotype is consistently observed across additional *MED13L* variant cell lines, we examined CCNC localization in the expanded fibroblast cohort using immunofluorescence microscopy. As expected, the only detectable CCNC in the control cell line was nuclear (Fig. 2A, top panels). However, aberrant CCNC cytoplasmic localization was observed in all *MED13L* variant cell lines (Fig. 2A, see Fig. S1-2 for additional images). These results indicate that aberrant CCNC nuclear release is a consistent feature across diverse *MED13L* genotypes.

**Figure 2.**
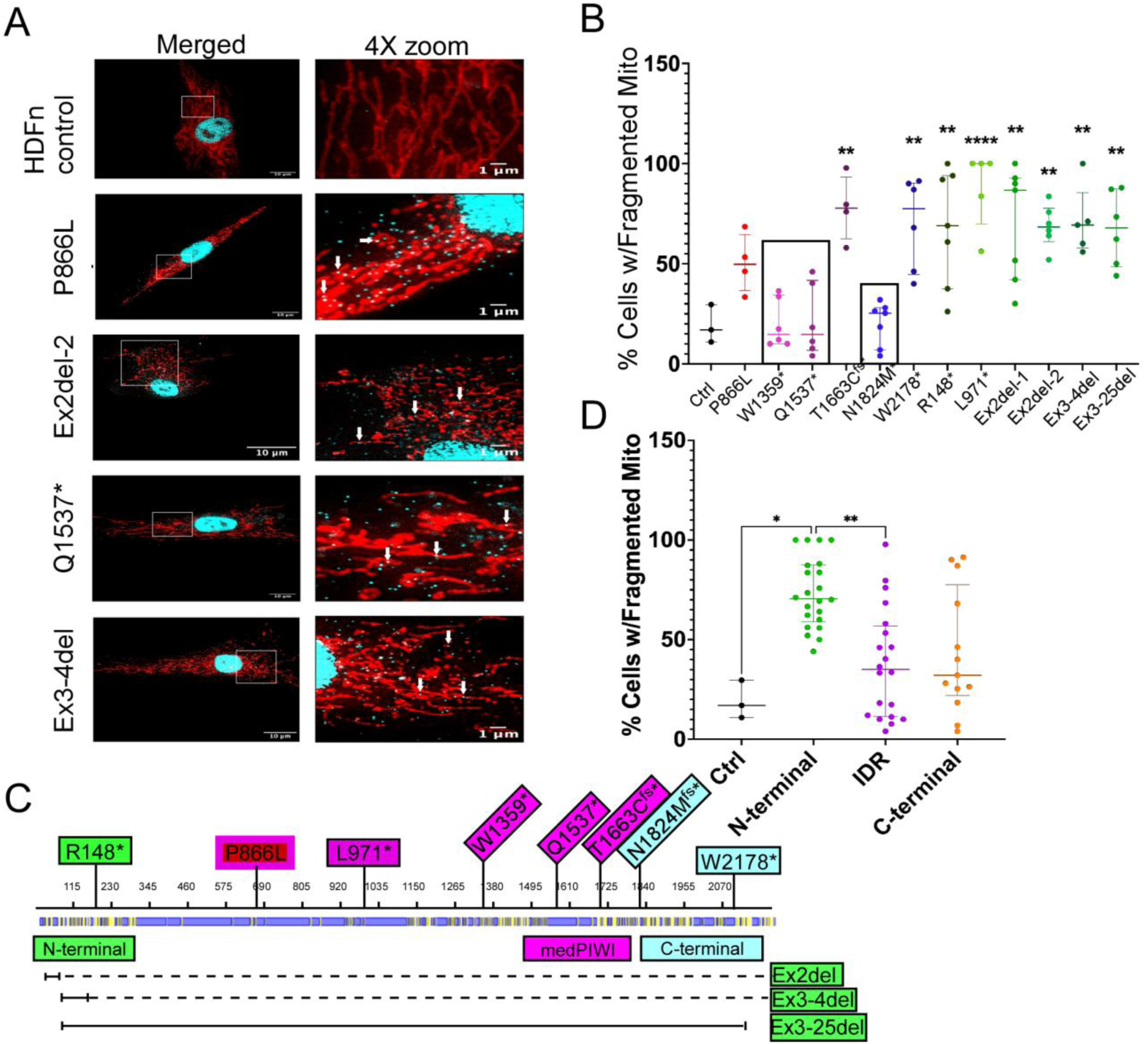
*MED13L* variants are associated with aberrant CCNC cytoplasmic localization and mitochondrial fragmentation. (A) Representative immunofluorescence images of mitochondrial morphology (red) and CCNC (cyan) in control and *MED13L*-variant fibroblasts as indicated. For each genotype, merged and 4x magnified images (indicated by boxes) are shown. White arrows indicate CCNC-mitochondrial overlap. See Figure S1-S2 for additional, full-field images. (B) Percentages of *MED13L* cells exhibiting >50% fragmented mitochondria are shown. Boxes highlight cell lines not displaying elevated mitochondrial fission. (n ≥ 100 cells analyzed). Control cells are represented in black, missense variant in red (P866L), medPIWI/IDR variants in shades of purple (L971*, W1359*, Q1537*, T1663C^fs*^), C-terminal variants in shades of blue (N1824M^fs^*, W2178*), and N-terminal or deletion variants in shades of green (R148*, Ex2del-1, Ex2del-2, Ex3-4del, Ex3-25del). Examples of mitochondrial fragmentation can be found in Figure S1. (C) UniProt-derived domain map and mutation locations of *MED13L* syndrome variants. Globular (yellow) and unstructured (blue) protein domains are indicated under the amino acid numbering. Frameshift, missense, and nonsense mutations are shown above the numbering, deletions (solid line) are indicated below. Dotted lines indicate predicted coding region eliminated by frameshift mutation in deletion alleles. The mutations were grouped based on position with N-terminal (green) mutations including one frameshift and three deletion alleles (R148*, Ex2del, Ex3-4del, Ex3-25del). The single missense variant (P866L) is represented by red box (variant type) with purple border (functional domain classification = IDR). IDR missense and truncation mutations, including those in the MID medPIWI domain, are represented with purple (L971*, W1359*, Q1537*, T1663C^fs*^). C-terminal mutations are represented in blue (N1823M^fs^*, W2178*). (D) Fragmented mitochondria phenotype was grouped by *MED13L* mutations as described in (C). All data represent mean ± SEM or distributions as indicated. Statistical comparisons were performed using one-way ANOVA with Dunnett’s post-hoc test comparing genotype or group to controls (*p < 0.05, **p < 0.01, *p < 0.001). See Table S1 for variant details.

We previously demonstrated that CCNC has been implicated in inducing mitochondrial fission through activation of the Dynamin-like GTPase DRP1^18,38^. We next assessed mitochondrial morphology in these cells using the stain MitoTracker Red^39^. As expected, the mitochondria in the control cell line appeared reticular with numerous branches (Fig. 2A, top panels). Compared with control cells, *MED13L*-mutant fibroblasts exhibited striking alterations in mitochondrial morphology, with many cell lines displaying short, punctuate mitochondrial fragments rather than elongated reticular networks (Fig. 2A, quantified in Fig. 2B). However, this phenotype was not uniformly observed across all cell lines. While the majority of *MED13L* variants exhibited fragmented mitochondria, several lines (W1359*, Q1537* and N1824M^fs^*, open boxes in Fig. 2B) displayed mitochondrial networks more comparable to control cells. These results indicate that fragmented mitochondria are a common, but not universal, feature associated with *MED13L* dysfunction. In addition, immunofluorescence analysis revealed increased CCNC signal overlapping mitochondrial networks in *MED13L* variant fibroblasts relative to controls (arrows, Fig. 2B). While this overlap does not demonstrate a direct molecular interaction, the redistribution of CCNC toward mitochondria is consistent with our previous reports linking stress-induced CCNC nuclear release to mitochondrial fragmentation and altered organelle dynamics.

One goal of this study was to determine whether mutation type or location can be predictive for *MED13L* syndrome outcomes. Therefore, *MED13L* mutations were mapped onto annotated protein domains, and grouped accordingly (Fig. 2C). Protein domains were based on known structural regions, as annotated in UniProt (MED13L_Human Q71F56). The frameshift, missense, and nonsense mutations are indicated on top of the map while the deletions are shown below. The solid line for the deletions represents the actual missing DNA while the dotted line indicates the truncation of the remainder of the protein due to the mutation. Using this mapping to create protein domain bins, this revealed domain-specific differences in mitochondrial morphology. N-terminal mutations (including all deletion mutations), which are predicted to truncate a substantial portion of the protein, exhibited consistently elevated mitochondrial fragmentation (Fig. 2D). Conversely, mutations within the intrinsically disordered regions (IDR) or C-terminal domains were not significantly different from control but did display variability within the group. These findings suggest that mutation location may contribute to variability in mitochondrial morphology, with N-terminal variants associated with more pronounced fragmentation. When considered alongside the consistent cytosolic CCNC localization, these data support the potential use of both mitochondrial fragmentation and CCNC mis-localization as candidate cellular biomarkers of *MED13L* syndrome.

### mtDNA depletion is a consistent biomarker of *MED13L*-associated mitochondrial dysfunction

Mitochondrial DNA (mtDNA) is organized into protein-DNA nucleoids, each containing 1–15 copies of the mitochondrial genome^40^. Previous studies have identified several molecules that maintain nucleoids including mtDNA binding proteins^41^ and factors controlling mitochondrial shape^42,43^. For example, constitutive mitochondrial fragmentation caused by deleting the fusion factors *MFN1* and *MFN2* resulted in nucleoid loss in individual mitochondria^44^. To assess whether *MED13L* variants impact mtDNA abundance and organization, we performed immunofluorescence microscopy using antibodies directed against dsDNA in conjunction with MitoTracker Red to observe morphology. Control fibroblasts exhibited dense, widely dispersed mtDNA foci throughout the mitochondrial network (Fig. 3A, left panel, overlapping signal shown in white). However, *MED13L* variant cell lines displayed visibly reduced mtDNA foci, with irregular distribution and regions of mitochondria lacking detectable nucleoids (Fig. 3A, arrow heads, right two panels).

**Figure 3.**
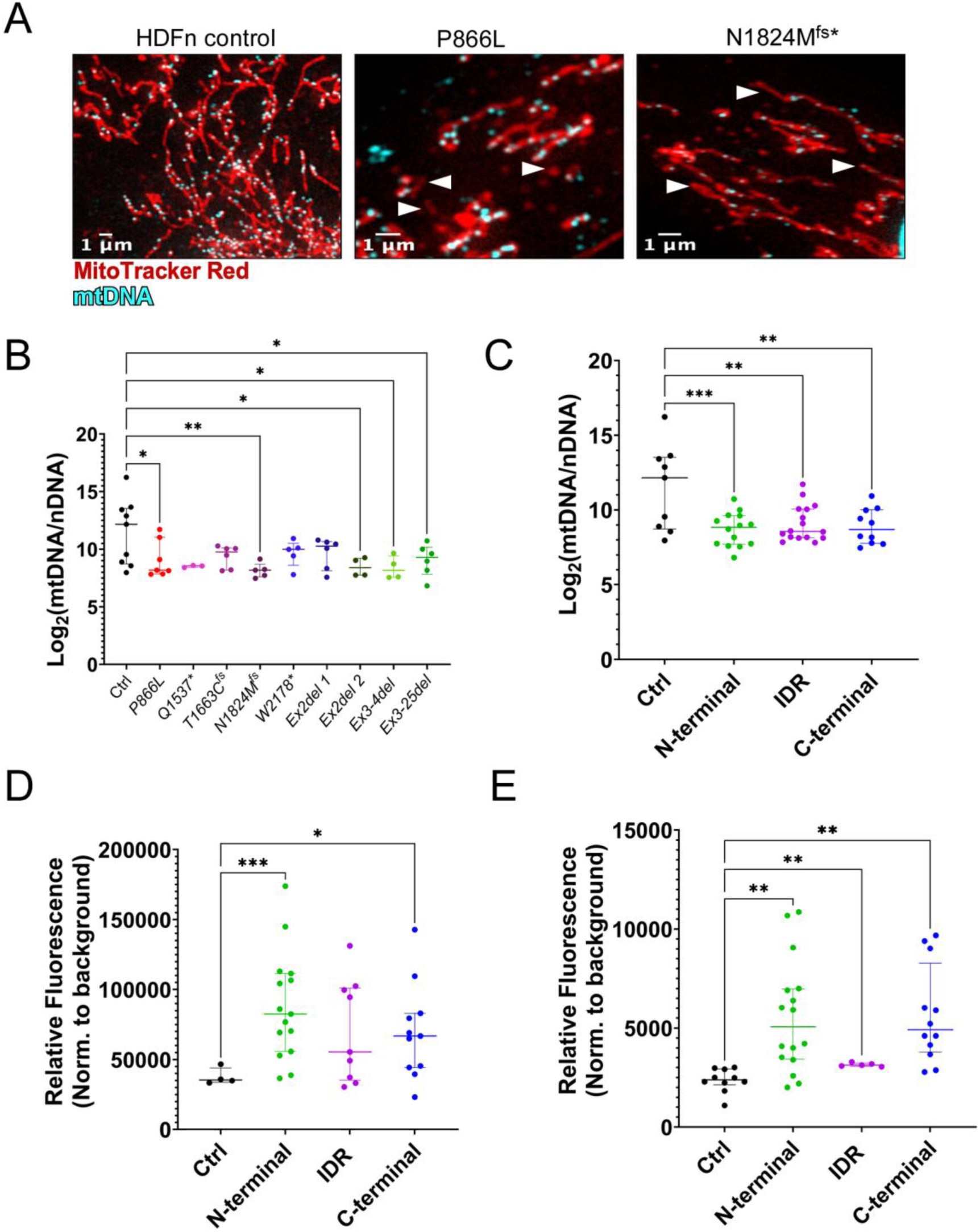
mtDNA depletion and increased oxidative stress are a consistent biomarker of *MED13L*-associated mitochondrial dysfunction. (A) Representative immunofluorescence images showing double-stranded DNA (dsDNA; blue) and mitochondria (red) in control and two *MED13L* variant fibroblasts (*P866L* and *N1824\**). Images depict mitochondrial network architecture and the distribution of mitochondrial nucleoids across genotypes. (B) Relative mitochondrial DNA (mtDNA) copy number in control and *MED13L* variant fibroblasts quantified by qPCR and normalized to the nuclear DNA standard (see methods) (n ≥ 3 biological replicates; technical triplicates). (C) mtDNA levels in control and mutations groups described in Fig. 2. (D) Cytosolic ROS levels measured using dihydroethidium (DHE) and presented as background-subtracted fluorescence values for control, missense, and PTV fibroblast lines (n ≥ 4 biological replicates; technical duplicates). (E) Quantification of mitochondrial ROS using MitoSOX fluorescence in control and *MED13L* variant fibroblasts classes described in Fig. 2. Relative fluorescence intensity (normalized to background) is shown (n ≥ 6 biological replicates; technical duplicate). All values represent mean ± SEM. Statistical comparisons were performed using one-way ANOVA with Dunnett’s post-hoc test. (*p < 0.05, **p < 0.01, ***p < 0.001, ****p < 0.0001).

To quantify these observations, the approximate mtDNA copy number was measured by qPCR^45,46^. Strikingly, all *MED13L* variant fibroblasts exhibited marked reductions in mtDNA content with many reaching statistical significance (Fig. 3B). Notably, these data revealed that mtDNA depletion represents a recurrent consequence of *MED13L* syndrome mitochondrial dysfunction. Next, we grouped variants according to protein domains, as before, to determine whether mtDNA copy number varied across domains. These results revealed a similar reduction in mtDNA content regardless of the mutation location (Fig. 3C). Taken together, these results indicate the loss of mtDNA content is a consistent phenotype associated with *MED13L* syndrome cells and is in parallel with reduced mitoATP production and increased organelle fragmentation.

Previous studies have revealed that extensive mitochondrial fragmentation often leads to elevated reactive oxygen species (ROS) due to disruption of the electron transport chain^47–49^. Therefore, we investigated whether the *MED13L* variant cell lines exhibited altered mitochondrial redox homeostasis. Using fluorophores dihydroethidium (DHE)^50^ and MitoSox Red^51^, we measured cytosolic and mitochondrial ROS levels, respectively. Binning the mutations according to protein domain revealed that both N- and C-terminal mutations resulted in elevated cytosolic ROS, with the N-terminal mutations exhibiting a stronger phenotype (Fig. 3D). For the IDR mutation, only the P866L cell line was examined, which trended upward but did not reach statistical significance. A similar result was obtained for mitochondrial ROS (Fig. 3E). Together, these findings identify reduced mtDNA abundance as a consistent feature of *MED13L* syndrome-associated mitochondrial dysfunction, accompanied by elevated oxidative stress in a subset of variants.

### *MED13L* variants display altered transcriptional dysregulation of MKM components and mitochondrial pathways

Prior transcriptional profiling in *Ccnc^-/-^*mouse embryonic fibroblasts (MEFs) revealed downregulation of genes required for mitochondrial maintenance and biogenesis^19,20^. In addition, our prior analysis of a single *MED13L* patient-derived fibroblast line similarly identified broad transcriptional changes affecting mitochondrial pathways^15^. To determine the extent to which *MED13L* variants are associated with transcriptional alterations, RT-qPCR analysis was performed on genes involved in mitochondrial biogenesis, metabolism, and MKM function. Across *MED13L* variant fibroblasts, we observe a consistent trend of transcriptional dysregulation. Surprisingly, we observed upregulation of other components of the MKM including *MED13*, *CCNC* and especially *MED13L* (Fig. 4A, detailed in 4B-D). These results suggest that compensatory activation of the other MKM genes occurs in a subset of *MED13L* variant cell lines. Although increased MKM transcript levels were observed, corresponding increases in protein abundance were not detected by Western blot analysis (Fig. S4). This discrepancy suggests the presence of post-transcriptional or translational regulatory mechanisms that modulate MKM protein levels in *MED13L* variants (Fig. S4A, B, C).

**Figure 4.**
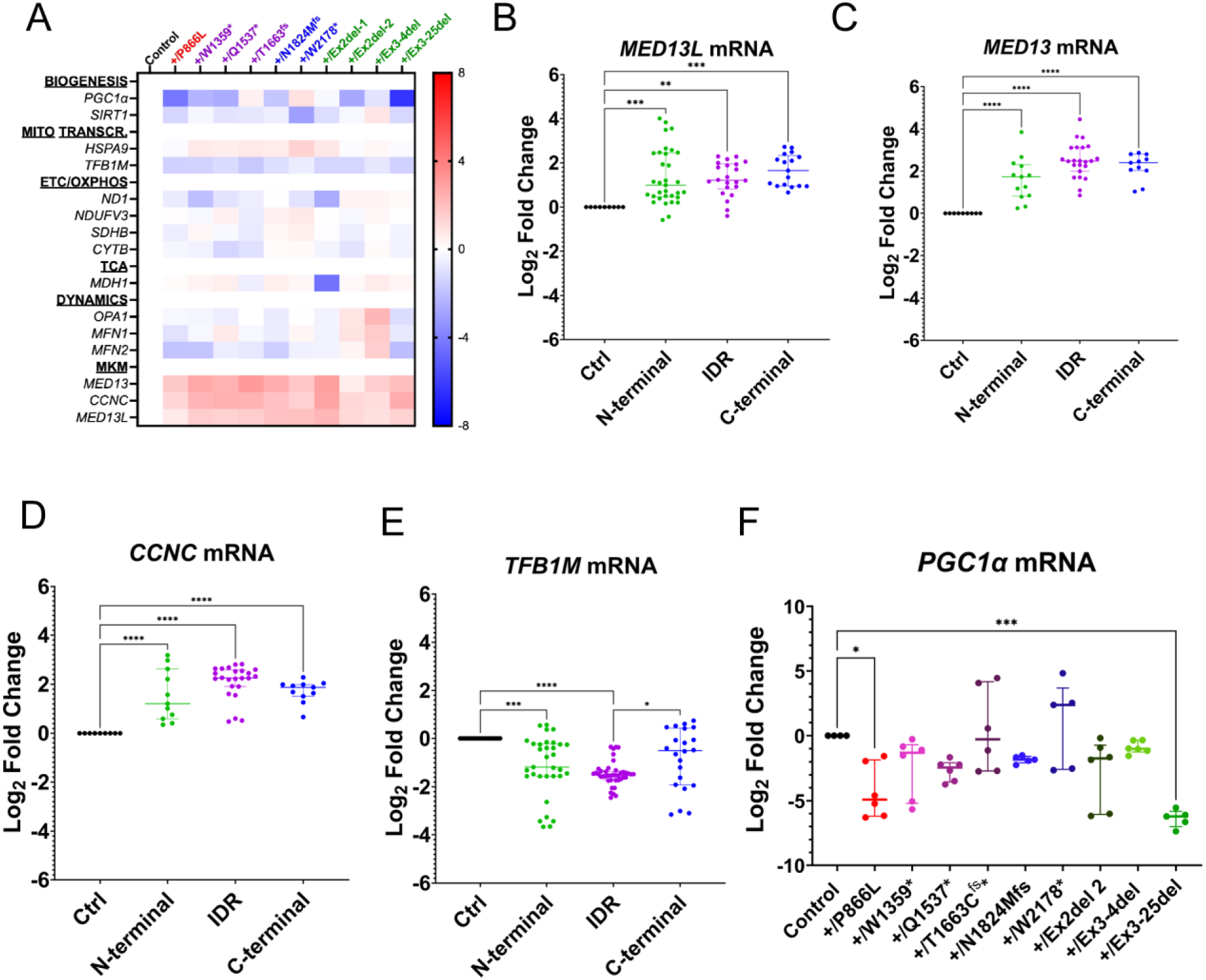
*MED13L* variants display altered transcriptional dysregulation of MKM components and mitochondrial pathways. (A) Heatmap of relative mRNA expression (log_2_FC vs. pooled controls) for genes involved in mitochondrial biogenesis (*PGC1ɑ, SIRT1*), mitochondrial transcription (*HSPA9, TFB1M*), electron transport chain and oxidative phosphorylation (*ND1, NDUFV3, SDHB, CYTB*), TCA cycle (*MDH1*), mitochondrial dynamics (*OPA1, MFN1, MFN2*), and Mediator Kinase Module components (*MED13, CCNC, MED13L*). Columns represent individual fibroblast lines stratified by mutation class as described in Fig. 2, and color-coordinated. Values are normalized to pooled control fibroblasts. Blue indicates relative downregulation and red indicates relative upregulation (n ≥ 4 biological replicates; technical triplicates). (B-F) Transcript RT-qPCR quantification for *MED13L*, *MED13*, *CCNC*, *TFB1M* and *PGC1ɑ* as indicated. All graphs are presented as log_2_ fold change relative to control fibroblasts. All RT-qPCR data represent median ± interquartile range; individual fibroblast lines plotted in heatmap. All data were assessed by one-way ANOVA with Dunnett’s post-hoc test comparing each mutation class to controls (*p < 0.05, **p < 0.01, ***p < 0.001, ****p < 0.0001).

Conversely, a subset of genes (*PGC1ɑ*^52^*, SIRT1*^53^*, TFB1M*^54^) involved in mitochondrial biogenesis and maintenance consistently displayed reduced mRNA levels in these cell lines. There was an overall reduction in *TFB1M* mRNA across cell lines tested, with C-terminal mutations showing a more modest phenotype compared to control (Fig. 4E). In addition, several cell lines including P866L, W1359*, Q1537*, N1824M^fs^*, Ex2del 2, and Ex3-25del displayed anywhere from a 1–8-fold reduction in *PGC1α* mRNA levels compared to control, with two of these reaching statistical significance (Fig. 4F). Interestingly, of these cell lines, the Q1537* and the C-terminal mutant N1824M^fs^* mutant cell lines exhibited normal mitochondrial morphology (Fig. 2B). These results suggest that their mitochondrial dysfunction may be the result of transcriptional, rather than a morphological, dysregulation

Together, these data reveal a conserved transcriptional signature across *MED13L* variant fibroblasts characterized by MKM component upregulation coupled with suppression of mitochondrial biogenesis and maintenance factors. Importantly, these results suggest the potential utility of gene expression as a molecular biomarker for *MED13L* variant-associated mitochondrial dysfunction.

### *MED13L* variants exhibit reduced proliferative capacity and features of cellular senescence

Senescence is defined by irreversible cell-cycle arrest coupled with an elevated inflammatory response known as the senescence-associated secretory phenotype (SASP)^55,56^. Although many SASP inducers are known, mitochondrial dysfunction is often observed in senescing cells, although the cause-and-effect relationship has not been completely established^55,57^. Hallmarks of senescence phenotypes include withdrawal from the cell cycle, expression of cyclin-dependent kinase inhibitors, and induction of senescence-related ß-galactosidase^58–60^. Given our finding of mitochondrial dysfunction, we next examined whether *MED13L* variant cells exhibited senescence phenotypes. Late-passage cultures corresponding to approximately 40-45 generations were assayed for proliferative capacity using BrdU incorporation assays^60^. Across the *MED13L* variants tested, the proportion of BrdU-positive cells was significantly reduced relative to controls (Fig. 5A), indicating decreased proliferative activity. These results indicate that a higher percentage of the *MED13L* variant population had withdrawn from mitotic cell division.

**Figure 5.**
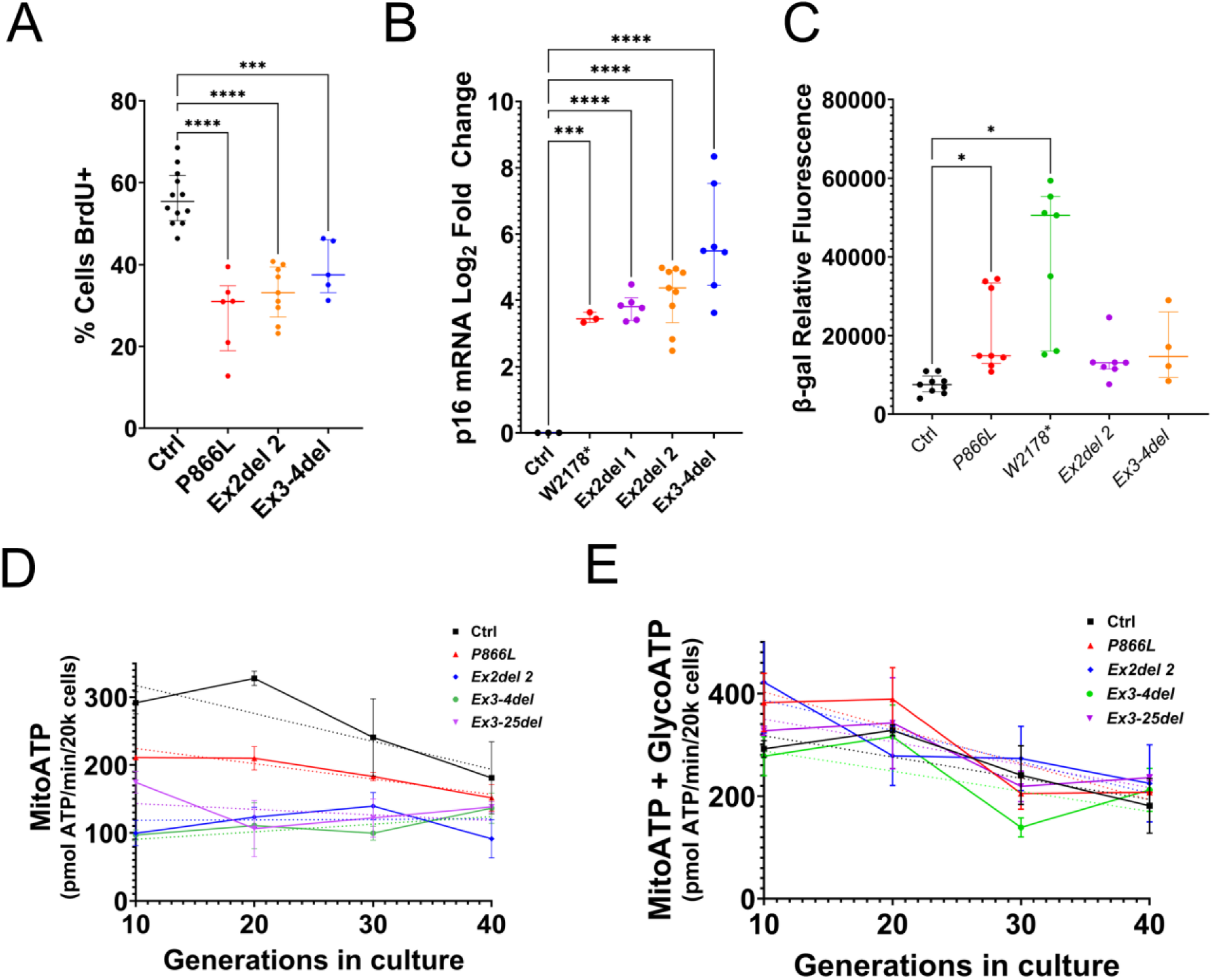
*MED13L* variants exhibit reduced proliferative capacity and features of cellular senescence. (A) Percentage of mitotically active cells determined by BrdU incorporation assays in control and representative *MED13L* variant fibroblasts. Reduced BrdU positivity indicates diminished proliferative capacity (n ≥ 5 biological replicates). (B) Relative expression of senescence marker p16^INK4a^ measured by RT-qPCR and normalized to *GAPDH*. Data are shown as Log_2_ fold change relative to control fibroblasts (n ≥ 3 biological replicates; technical triplicates). (C) Senescence-associated β-galactosidase (SA-β-gal) activity quantified by flow cytometry using a fluorogenic β-gal substrate. Background-subtracted fluorescence intensity is shown for control and select *MED13L* variant fibroblasts (n ≥ 4 biological replicates). (D, E) MitoATP production (D) or total ATP production (E) in extended cell cultures for control and representative *MED13L* variant fibroblast lines. Timepoints taken every 10 generations. Simple linear regression and best fit analysis performed, as shown by indicated corresponding dotted lines. Data represent individual biological replicates with summary statistics displayed as median and interquartile range. Statistical analysis was performed using one-way ANOVA with Dunnett’s post hoc correction for variant-to-control comparisons. (*p < 0.05, **p < 0.01, ***p < 0.001, ****p < 0.0001).

To further interrogate a potential elevated senescence entry, two additional assays were performed. First, we used RT-qPCR to measure the expression of the cyclin-dependent kinase inhibitor p16 (*CDKN2A*), a canonical marker of irreversible growth arrest^61–63^. Four of the five variants examined displayed statistically significant elevation in p16 mRNA compared to control (Fig. 5B). Finally, we quantified senescence-associated β-galactosidase activity using a fluorogenic flow cytometry-based assay^64,65^. These studies revealed mixed results with P866L and W2178* cell lines expressing elevated ß-galactosidase, while the two deletion alleles did not (Fig. 5C). Together, these data suggest that late passage *MED13L* variant fibroblasts exhibit reduced proliferative capacity and features consistent with senescence, although individual senescence markers are variably expressed across cell lines. Our finding that some cell lines did not express all senescence markers is a common issue for these types of studies^59^. These findings suggest that mitochondrial dysfunction in *MED13L* syndrome may not simply be a static metabolic defect but may actively contribute to cellular aging and loss of tissue homeostasis.

As mitochondrial dysfunction and senescence are closely linked, we next examined whether mitochondrial energetic capacity changed during prolonged culture. Mitochondrial ATP production was therefore assessed across serial passage in control and *MED13L* fibroblasts. Three independent cultures from four *MED13L* variant fibroblast lines and controls were continuously maintained for approximately 40 generations, with mitoATP measurements obtained every ten generations (Fig. 5D). At the earliest analyzed timepoint (generation 10), *MED13L* fibroblasts already exhibited reduced mitochondrial ATP production relative to control cells, consistent with our earlier observations. As cultures continued to age in vitro, mitoATP production progressively declined across all fibroblast lines, including the control. However, two-way ANOVA identified a significant interaction between fibroblast line and generation number (P = 0.0011), indicating that longitudinal mitoATP trajectories differed between genotypes during serial passaging. A significant overall effect of fibroblast line (genotype) was additionally observed (P < 0.0001), whereas generation number alone did not significantly contribute to overall variance (time). Linear regression analysis further demonstrated distinct mitoATP trajectories between fibroblast lines, with control fibroblasts exhibiting the steepest negative slope across serial generations. In contrast, several *MED13L* fibroblast lines maintained persistently reduced mitochondrial ATP production beginning in early generations with comparatively limited longitudinal change. By later generations, mitoATP production converged toward similarly reduced levels across all lines, suggesting that *MED13L* disruption may promote an early-established and sustained bioenergetic state resembling later-passage control fibroblasts.

Analysis of total ATP production revealed a similar overall pattern, with *MED13L* variant fibroblasts exhibiting persistently impaired energetic output throughout serial passaging (Fig. 5E). While certain fibroblast lines maintained relatively stable but chronically reduced ATP levels, others demonstrated more progressive declines across culture duration (P866L, red). Together, these findings indicate that *MED13L* disruption is associated with senescence-like cellular remodeling characterized by reduced proliferative capacity, elevated p16 expression, and persistent bioenergetic dysfunction during prolonged culture.

### Mitochondrial function is associated with mutation position, participant age, and clinical phenotypes

The results presented in this study indicate that the *MED13L* variant cell lines exhibit several phenotypes that vary in severity and scope. For example, although mitochondrial activity is reduced in all cell lines examined, the extent of these deficiencies varies, typical of a spectral disorder. We next tested whether the mutation location affected mitochondrial function and could thereby be predictive for untested mutant alleles. To investigate these possibilities, we examined mitoATP production using the previously described mutation location categories. These results revealed that the N-terminal mutations, which consist mostly of deletions that cause early truncation alleles, exhibited the most severe reduction in ATP production (Fig. 6A). The IDR/medPIWI domain mutations were the next most severe, while the C-terminal alleles were the least affected and were statistically different than the N-terminal mutant cell lines. These results suggest a positional effect, with N-terminal variants associated with more severe impairment of mitochondrial function.

**Figure 6.**
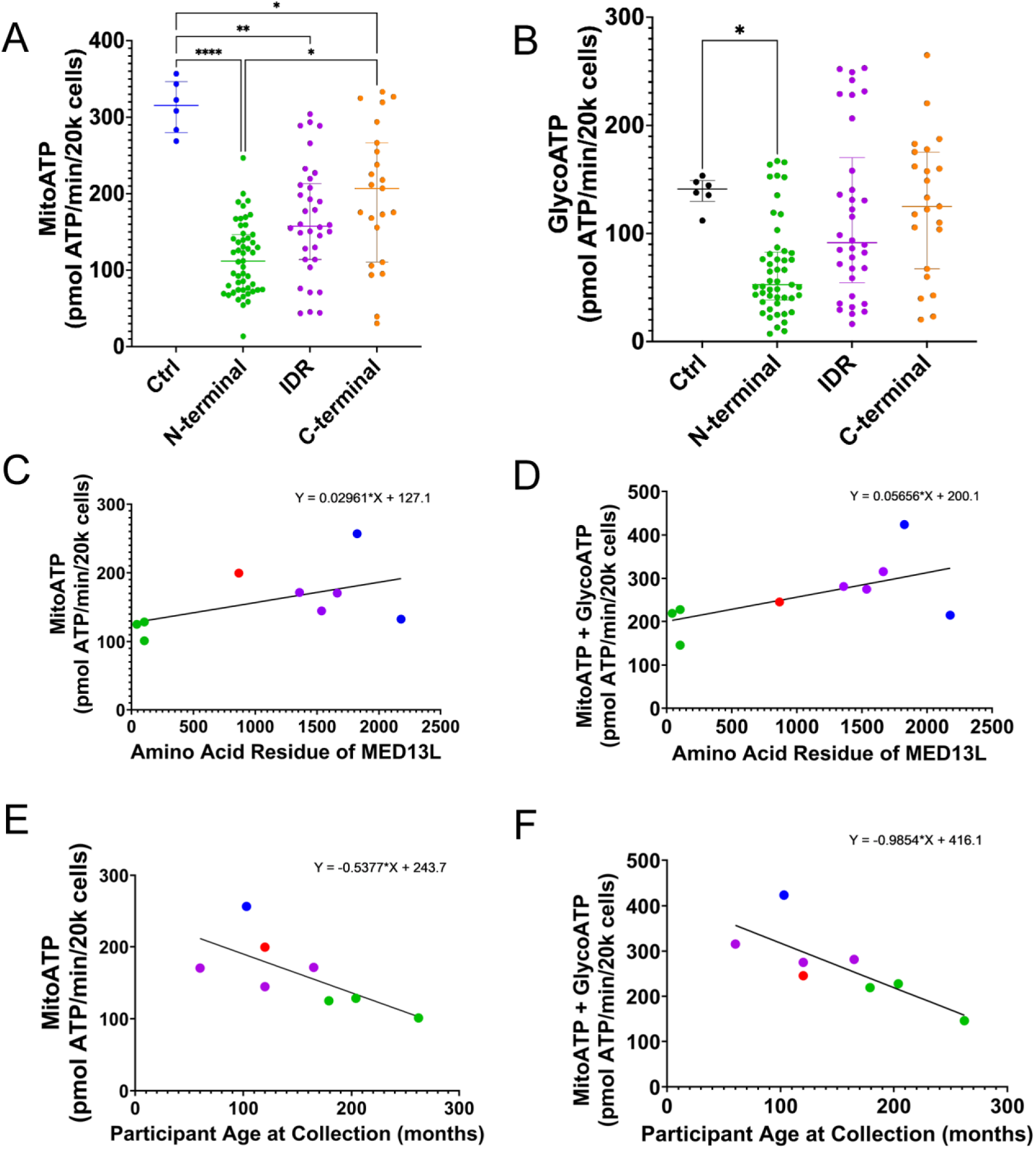
Mitochondrial function is associated with mutation position, participant age, and clinical phenotypes. (A) Mitochondrial ATP production (mitoATP) measured in control and *MED13L* patient-derived fibroblasts grouped by variant location (IDR, N-terminal, MedPIWI or C-terminal). Each point represents an independent biological replicate (n ≥ 6). (B) Glycolytic ATP production (glycoATP), calculated from proton efflux rates, shown by variant class as in (A) (n ≥ 6). (C) Relationship between mitoATP production (mean values from Fig. 1) and MED13L amino acid position of each variant. Each point represents a fibroblast line carrying a variant at the indicated residue position within the MED13L protein (R = 0.515). (D) Relationship between total ATP production (mitoATP + glycoATP) and MED13L amino acid position of mutation. Each point represents an individual fibroblast line carrying variants at the indicated residue position. (R = 0.597) (E) Relationship between mitoATP production and participant age at time of sample collection, in months (R = 0.697). (F) Relationship between total ATP production (mitoATP + glycoATP) and participant age at time of sample collection, in months (R = 0.775). Lines in (C-F) represent line of best fit following simple linear regression. Corresponding equations are represented on each graph. For (C-F), color of the symbol corresponds to type and/or location of mutation and are as follows: green = N-terminal deletions, red = missense, purple = IDR/medPIWI nonsense and frameshift, blue = C-terminal nonsense and frameshift.

To determine whether glycolytic compensation is different across variant classes, glycolytic ATP production (glycoATP) was also assessed (Fig. 6B). In contrast to mitoATP production, glycoATP measurements demonstrated substantial variability between fibroblast lines and did not exhibit a consistent potential trend. Several variant lines displayed elevated glycolytic ATP production relative to controls, suggesting partial metabolic compensation for impaired mitochondrial energetic capacity in select fibroblast populations.

We next directly assessed the relationship between mutation position and energetic function using simple linear regression analyses. MitoATP production demonstrated a positive relationship with the position of the *MED13L* amino acid mutation, with more C-terminal variants generally exhibiting higher mitochondrial ATP production relative to variants located nearer the amino terminus of the protein (R = 0.515; Fig. 6C). A similar trend was observed for total ATP production, in which more carboxy-terminal mutations were associated with higher overall energetic output (R = 0.597; Fig. 6D). Together, these findings suggest that variants occurring earlier within the *MED13L* coding sequence are associated with more severe mitochondrial dysfunction.

As clinical severity in neurodevelopmental disorders may also change with age, we next examined whether mitochondrial energetic capacity correlated with participant age at the time of fibroblast collection. In contrast to mutation position, mitoATP production displayed a negative relationship with participant age (R = 0.697; Fig. 6E), indicating that fibroblasts derived from older participants generally exhibited lower mitochondrial ATP production. Total ATP production similarly declined with increasing participant age (R = 0.775; Fig. 6F). Although the cohort size remains limited, these findings raise the possibility that mitochondrial dysfunction in *MED13L* syndrome may worsen over time or reflect cumulative age-associated bioenergetic stress. Collectively, these data demonstrate that mitochondrial dysfunction in *MED13L* syndrome is heterogeneous and influenced by both mutation position and participant age, with earlier variants and older participant-derived fibroblasts generally exhibiting more severe energetic impairment.

### Clinical and caregiver-reported functional measures are associated with *MED13L* mutation location

To contextualize our molecular analyses, we first summarized clinician-rated and caregiver-reported functional outcomes across multiple standardized assessment tools (see methods). First, we examined motor ability scores using the Vineland Adaptive Behavior Scales^31,32,66,67^ incorporating both fine and gross motor skills. These raw scores were plotted against the amino acid location of the *MED13L* mutation (Fig. 7A). This analysis revealed a trend similar to mitoATP production in that the N-terminal mutations exhibited lower motor function compared to those with more C-terminal mutations. In contrast, no clear relationship was observed between motor scores and patient age within the available dataset (Fig. S7), although the number of individuals with complete data was limited.

**Figure 7.**
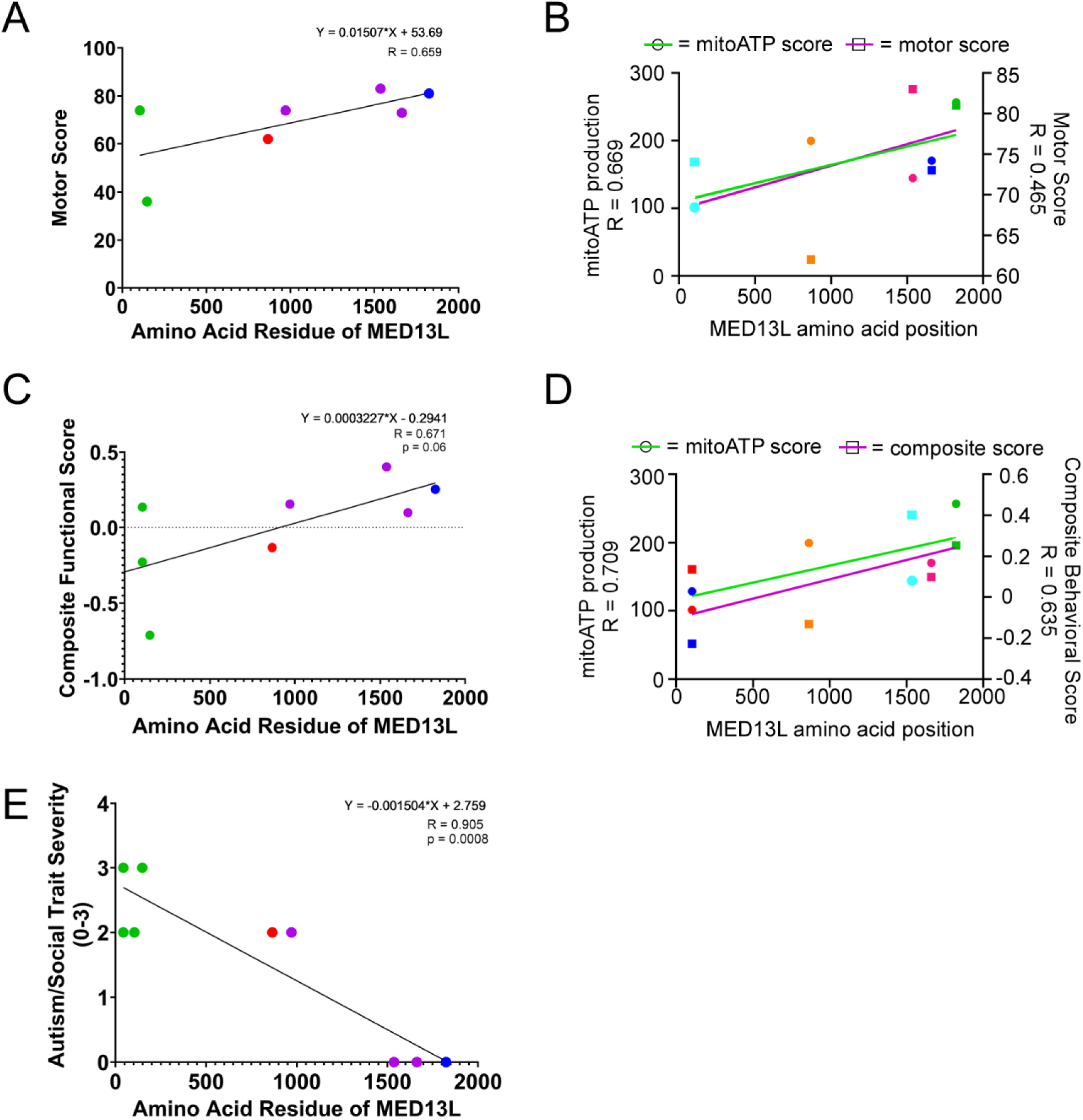
Clinical and caregiver-reported functional measures associate with *MED13L* mutation location. (A) Vineland motor standard scores (combined gross and fine motor domains) plotted against MED13L amino acid position. Each point represents individual participant harboring indicated *MED13L* variant. Linear regression performed, best fit line shown (R = 0.659). (B) Dual axis comparison of Vineland motor severity scores (R = 0.465; magenta) with mitochondrial ATP (R = 0.669; mitoATP; green) and amino acid location of the variant. Independent linear regression analyses were performed for each variable relative to motor severity. (C) Composite functional severity scores generated from integrated adaptive, communication, and motor assessments (see Methods) plotted against MED13L amino acid position. Linear regression and line of best fit are shown (R = 0.671, p = 0.06) (D) Dual-axis comparison of composite functional severity scores (R = 0.635; magenta line) with mitochondrial ATP production (R = 0.709; mitoATP; green) and MED13L amino acid position. Independent linear regression analyses were performed for each variable relative to composite severity score. (E) Autism/social trait severity scores generated from harmonized caregiver-reported and clinical behavioral datasets plotted against MED13L amino acid position. Linear regression and line of best fit are shown (R = 0.905, p = 0.0008). For all panels, amino acid position corresponds to the predicted location of the MED13L protein alteration. Each point represents an individual participant-derived fibroblast line. For (A, C, and E), color of the symbol corresponds to type and/or location of mutation and are as follows: green = N-terminal deletions, red = missense, purple = IDR/medPIWI nonsense and frameshift, blue = C-terminal nonsense and frameshift.

To further evaluate the relationship between clinical severity and mitochondrial dysfunction, mitochondrial ATP production and mutation position were simultaneously plotted against motor severity scores using dual y-axes (Fig. 7B). Independent linear regression analyses demonstrated that reduced mitoATP production was associated with increased motor severity, with earlier variant positions similarly trending toward greater impairment. The parallel regression patterns support a relationship between mutation position, mitochondrial dysfunction, and clinical motor outcomes in *MED13L* syndrome.

Composite functional/behavioral severity scores, generated from integrated adaptive, motor, and communication outcome measures (as in Fig. 6; see methods for details), demonstrated a similar positional trend, with more amino terminal *MED13L* variants associated with greater overall behavioral/functional dysfunction (Fig. 7C). To further assess the relationship between composite scores and mitochondrial dysfunction, mitoATP production and MED13L amino acid mutation position were simultaneously plotted against composite scores using dual axes as in Fig. 7B. Independent regression analyses demonstrated that lower mitochondrial ATP production and earlier variant positions were both associated with increased composite behavioral and functional severity, further linking impaired cellular bioenergetics with multisystem clinical dysfunction.

Finally, autism trait severity revealed an inverse correlation with mutation location. Variants located toward the N-terminal region were associated with higher autism/social trait severity scores, whereas variants positioned more distally within the protein were generally associated with milder social-behavioral phenotypes (Fig. 7E). Taken together, these results suggest that mutation location and mitochondrial ATP production are associated with clinical variability in *MED13L* syndrome. While these relationships should be interpreted cautiously given the cohort size, they support a model in which genotype and cellular bioenergetic status contribute to functional outcomes in affected individuals.

## Discussion

Prior *MED13L* syndrome studies have been limited by small cohorts, heterogeneous clinical presentations and a lack of mechanistic convergence. Here, using the largest panel of patient-derived *MED13L* fibroblasts analyzed to date, we identify mitochondrial impairment as a consistent and distinguishing cellular signature across diverse pathogenic variants. Despite substantial heterogeneity in transcriptional and morphological outcomes, all *MED13L* mutant fibroblast lines exhibited reduced mitochondrial content, diminished ATP production, and marked loss of mtDNA copy number, supporting mitochondrial impairment as a central aspect of *MED13L*-associated cellular dysfunction. Among the cellular phenotypes examined, reduced mitochondrial ATP production emerged as the most reproducible feature across genotypes, suggesting that impaired oxidative bioenergetics may represent a core downstream consequence of *MED13L* syndrome dysfunction.

These findings are consistent with growing evidence linking mitochondrial dysfunction to neurodevelopmental and neurodegenerative disorders, including conditions characterized by impaired oxidative phosphorylation, reduced mtDNA copy number, and elevated oxidative stress^35,37,68,69^. In this context, the increased cytosolic and mitochondrial ROS observed in a subset of *MED13L* variant cell lines further supports disruption of mitochondrial homeostasis as a contributing feature of disease pathology. Enhanced oxidative stress is associated with several intellectual disability disorders including autism^37,70–74^ as well as several neurological syndromes including Friedreich’s ataxia^75–77^ and Parkinson’s disease^78–80^. Neurons may be especially vulnerable to disruptions in mitochondrial homeostasis because of their high energetic demand, dependence on mitochondrial trafficking, and limited glycolytic reserve. Therefore, the role for mitochondrial dysfunction and elevated oxidative stress observed across *MED13L* variant fibroblasts is consistent with pathologies identified for other neurological disorders and supports the possibility that impaired mitochondrial resilience contributes directly to neurodevelopmental dysfunction seen in *MED13L* syndrome.

A key observation in this study is the consistent cytoplasmic localization of cyclin C (CCNC) across all *MED13L* variant cell lines. CCNC nuclear release has been previously implicated in stress-responsive mitochondrial fission through interaction with DRP1, suggesting a potential link between MKM dysfunction and altered mitochondrial dynamics^14,18,38,81^. Mitochondrial dynamics are controlled by the opposing activities of mitochondrial fusion (MFN2 and OPA1) and fission (DRP1) factors^82–92^. Disruption of this process, such as with *OPA1* mutations, results in excessive mitochondrial fragmentation in neurodegenerative diseases such as Charcot-Marie Tooth type 2A and Kjer disease/autosomal dominant optic atrophy^93–98^. Combined with our earlier demonstration of respiratory defects in *MED13L* fibroblasts, these findings support a model in which MKM dysfunction impairs nuclear regulation of mitochondrial homeostasis as well. While our data does not directly establish causality, the co-occurrence of CCNC mis-localization and mitochondrial fragmentation in many variant lines supports a model in which disruption of MKM-associated signaling contributes to altered mitochondrial structure and function.

Importantly, not all mitochondrial-associated phenotypes were uniformly observed across variant lines. While reduced mitoATP production and mtDNA depletion were highly consistent, mitochondrial morphology, ROS accumulation, glycolytic compensation, and senescence-associated markers displayed greater variability across the cohort. This heterogeneity likely reflects differential effects of variant position, residual protein function, compensatory metabolic adaptation, and the patient-specific genetic background. Nonetheless, the persistence of mitochondrial energetic dysfunction across diverse mutation classes suggests convergence upon a shared downstream cellular phenotype despite substantial molecular and clinical variability.

Our transcriptional analyses further support disruption of coordinated mitochondrial regulatory programs in *MED13L* syndrome. Previous studies of *Ccnc*-deficient mouse embryonic fibroblasts (MEFs) identified altered expression of genes involved in mitochondrial maintenance and biogenesis^19,20^. Consistent with these findings, *MED13L* variant fibroblasts exhibited reduced expression of multiple genes associated with mitochondrial function, including *PGC1α*, *SIRT1*, and *TFB1M*. Surprisingly, we also observed increased expression of several MKM-associated transcripts across many variant lines, suggesting compensatory transcriptional activation in response to impaired MKM function. However, corresponding increases in MED13L protein abundance were not detected by Western blot analysis, indicating that elevated transcript levels alone may be insufficient to restore MKM activity (Fig. S4A-B, quantified in S4C). Finally, additional Western blot analysis failed to detect truncated intermediates in whole cell extracts (Fig. S4E). Although a negative result, these findings suggest that truncation and deletion mutations both rendered cells haplo-insufficient. Conversely, this may suggest that truncated proteins are present at levels below the limits of detection. Regardless, this discordance further supports the possibility that multiple *MED13L* variants ultimately function through haploinsufficiency or impaired protein stability despite apparent transcriptional compensation. These findings additionally suggest that therapeutic strategies aimed solely at increasing *MED13L* transcript abundance may not fully restore MKM function due to failure to recover normal protein expression.

An additional finding of this study is the possibility that mitochondrial dysfunction in *MED13L* syndrome may represent a progressive rather than static phenotype. Longitudinal analysis revealed a persistent impairment of mitochondrial ATP production during extended culture, indicating that *MED13L* variants compromise metabolic resilience rather than immediately disabling respiration. This progressive vulnerability coincided with premature acquisition of senescence-associated phenotypes, including growth arrest, p16 induction, and accumulation of β-galactosidase-positive cells^57,59,62,64,102–106^. Notably, several *MED13L* fibroblast lines exhibited persistently reduced mitoATP production beginning at early timepoints, suggesting that *MED13L* disruption may establish an early and sustained bioenergetic state resembling later-passage control fibroblasts. These data support a model in which *MED13L* mutation accelerates cellular aging through cumulative mitochondrial stress, rather than causing acute toxicity. Although these findings were observed in vitro, this temporal pattern may help explain phenotypic evolution in patients, in whom early developmental delay can be followed by worsening neurological, metabolic, or systemic features over time, as reported by many parents and clinicians (caregiver-reported regression).

Importantly, our findings also suggest that mutation position may partially influence mitochondrial and clinical severity. Earlier truncating and deletion variants were generally associated with greater reductions in mitochondrial ATP production, while more C-terminal variants displayed comparatively preserved energetic function. Similar positional relationships were observed for motor function, composite functional and behavioral severity, and autism/social trait scores. Although these analyses remain limited by cohort size, they raise the possibility that disruption of distinct MED13L structural regions differentially impacts MKM stability and downstream mitochondrial regulation. More broadly, these findings support the concept that mutation position may contribute to phenotypic stratification in *MED13L* syndrome and may help predict cellular bioenergetic dysfunction.

Finally, these results have broader implications for the Mediator Kinase Module and related neurodevelopmental disorders. *MED12*^107–111^, *MED12L*^112^*, MED13*^113–115^, *CDK8*^116–118^, and *CDK19*^119–121^ mutations are linked to phenotypes similar to *MED13L* syndrome, including developmental delays and intellectual disability^122^. Due to these mutations producing overlapping neurodevelopmental phenotypes, this raises the possibility that mitochondrial dysfunction may represent a broader downstream consequence of MKM disruption. Whether CCNC itself contributes similarly remains unclear, as there have currently been no reports of mutations to this gene. Although constitutive CCNC nuclear release promotes mitochondrial fragmentation, heterozygous *Ccnc* loss in murine models does not recapitulate severe developmental phenotypes, suggesting that aberrant CCNC localization, rather than simple dosage reduction, may be critical for mitochondrial dysfunction^38,123^.

In summary, our work identifies mitochondrial dysfunction as a consistent and quantifiable biomarker of *MED13L* syndrome. These findings establish a mechanistic foundation for linking Mediator kinase module dysfunction to impaired mitochondrial homeostasis and provide a framework for therapeutic discovery aimed at restoring mitochondrial integrity. In addition, we find that mutation location approximates mitochondrial function suggesting that identifying the DNA lesion can be predictive of mitochondrial activity at least in fibroblast cell lines. Several limitations of the study should be considered. This study is based on fibroblast models, which may not fully recapitulate tissue-specific effects in neuronal or cardiac systems^124^. Further analysis in neuronal cell models would help establish these phenotypes. Despite fibroblasts not fully recapitulating neuronal biology, patient-derived fibroblast have proven valuable for identifying conserved mitochondrial and metabolic phenotypes across many neurological disorders including Alzheimer’s disese^125–131^. In addition, the cohort size limits the ability to definitively resolve the relative contributions of mutation type, position, and patient-specific factors such as age. Future studies incorporating larger cohorts and disease-relevant cell types, including neuronal models, will be important for validating and extending these findings.

Overall, this work identifies mitochondrial dysfunction as a convergent and quantifiable cellular feature of *MED13L* syndrome and establishes a functional connection between Mediator kinase module disruption and impaired mitochondrial homeostasis. By defining shared cellular phenotypes across *MED13L* variants, this study advances the field beyond genotype-based classification and toward a biomarker-informed understanding of Mediator-associated neurodevelopmental diseases. These findings provide a framework for future therapeutic strategies aimed at restoring mitochondrial integrity and improving cellular resilience in *MED13L* syndrome and related MKM-associated disorders.

## Acknowledgments

We would like to thank the MED13L Foundation for the organization of collection of patient-derived samples, in conjunction with CUIMC. We would also like to thank all of the families and affected individuals for volunteering their time and samples to complete such a large characterization of *MED13L* syndrome. We thank C. Greuter (University of Iowa) for helpful comments during this study.

## AUTHOR CONTRIBUTIONS

A.N.C.: Conceptualization, methodology, formal analysis, investigation, resources, writing-original draft, writing-review and editing, visualization, supervision, project administration, funding acquisition

K.N.J.: Validation, investigation, methodology, formal analysis

S.J.D.: Validation, formal analysis, investigation, writing-review and editing

J.M.B.: Methodology, validation, investigation, resources, project administration

E.L., B.C., J.F., and C.L.R.: Validation, investigation, methodology, resources, writing-review and editing

R.S.: Conceptualization, validation, resources, writing-original draft, writing-review and editing, visualization, supervision, project administration, funding acquisition

