## Supplemental Figures & Tables for "Cyclin C nuclear release and mitochondrial dysfunction define molecular signatures of *MED13L* Syndrome"

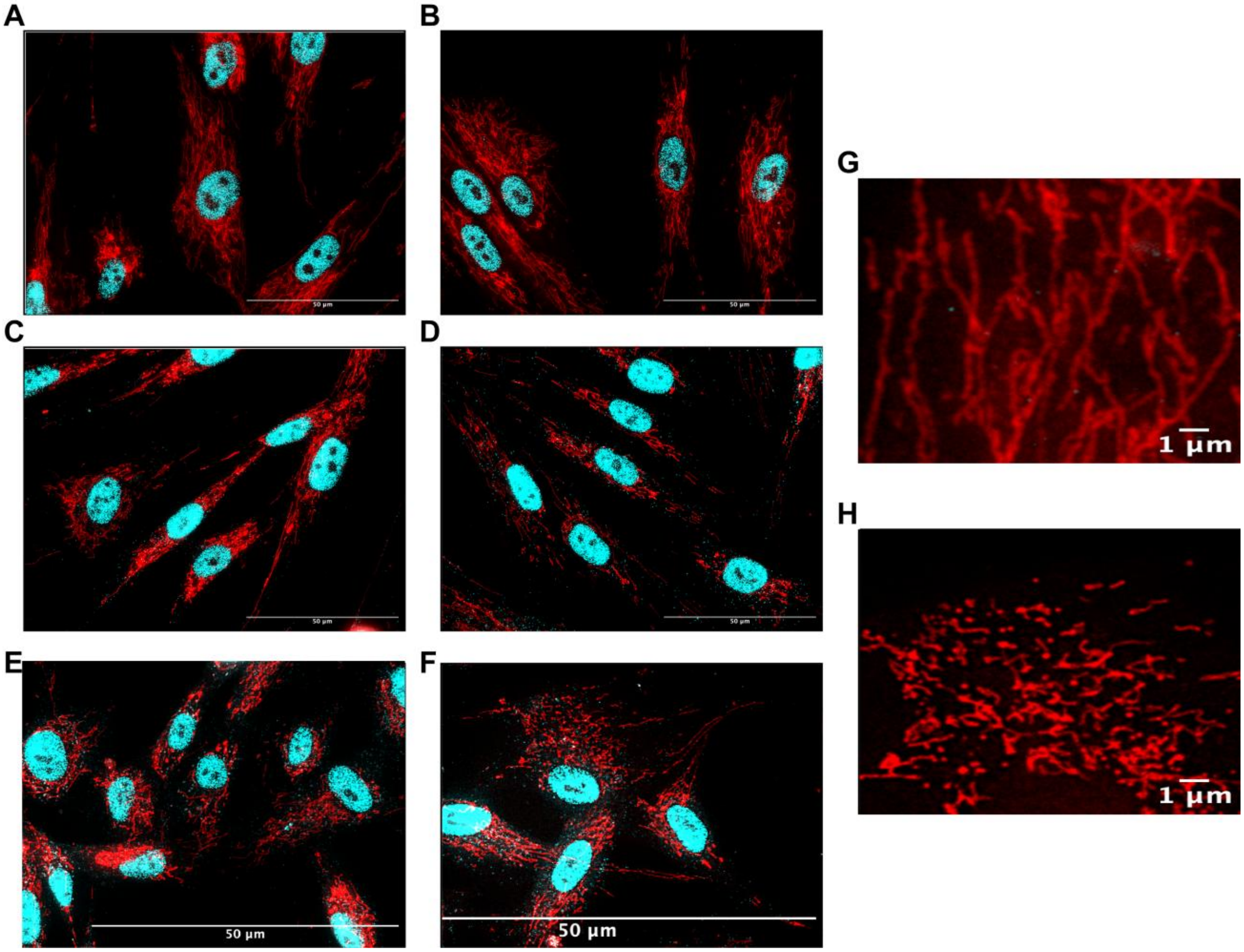


**Figure S1. Full-field immunofluorescence images of mitochondrial morphology and CCNC localization.** (A-F) Representative full-field immunofluorescence microscopy images of HDFn control (A + B), P866L (C + D), and Ex2del (E + F) patient fibroblasts corresponding to the cropped panels shown in Figure 2. (G-H) Representative examples of mitochondrial network morphologies used to define scoring criteria for mitochondrial fragmentation analysis. Cells displaying an elongated, interconnected mitochondrial network were classified as fused (G), whereas cells with numerous short, punctate mitochondria lacking network connectivity were classified as fragmented (H). These examples illustrate the morphological extremes used to guide phenotypic classification during quantitative analysis.

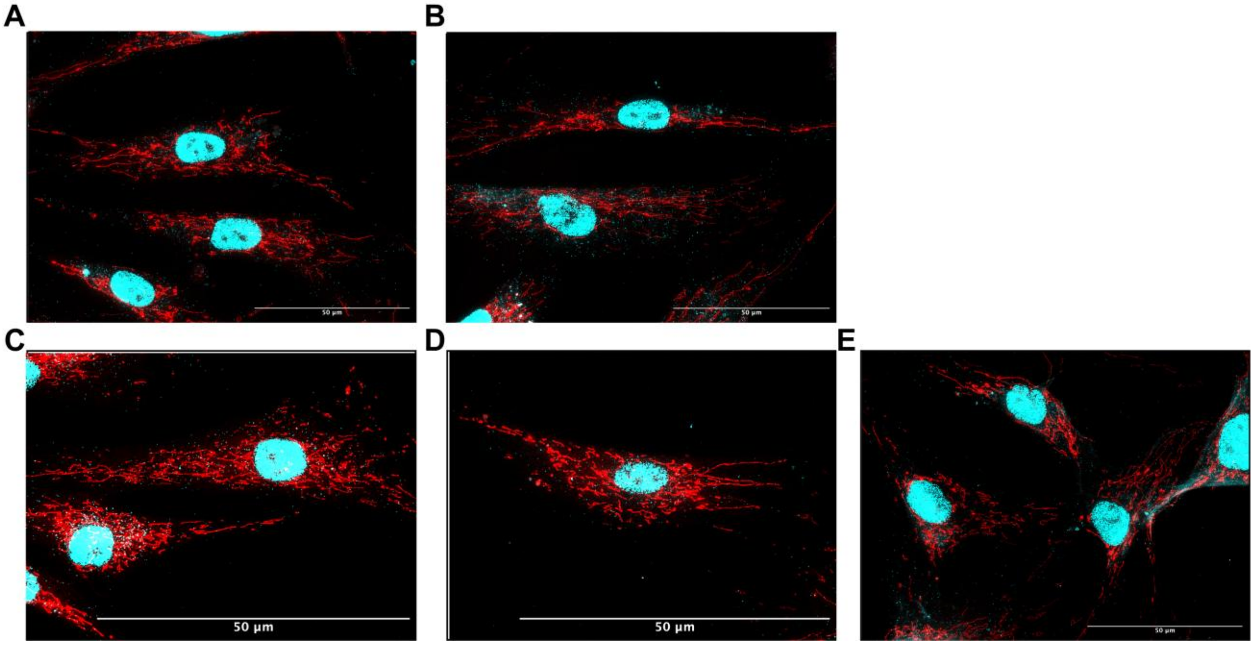


**Figure S2. Full-field immunofluorescence images corresponding to cropped panels shown in Figure 2.** Representative full-field microscopy images of *MED13L* patient samples harboring Q1537* (A-B) and Ex3-4del (C-E). These images represent the uncropped fields of view from which the regions shown in the main figure were selected, as well as additional full-field views as needed (B, D, E). Full-field images are provided to demonstrate the overall staining distribution and to confirm that representative regions were chosen for presentation. Scale bars, 50um

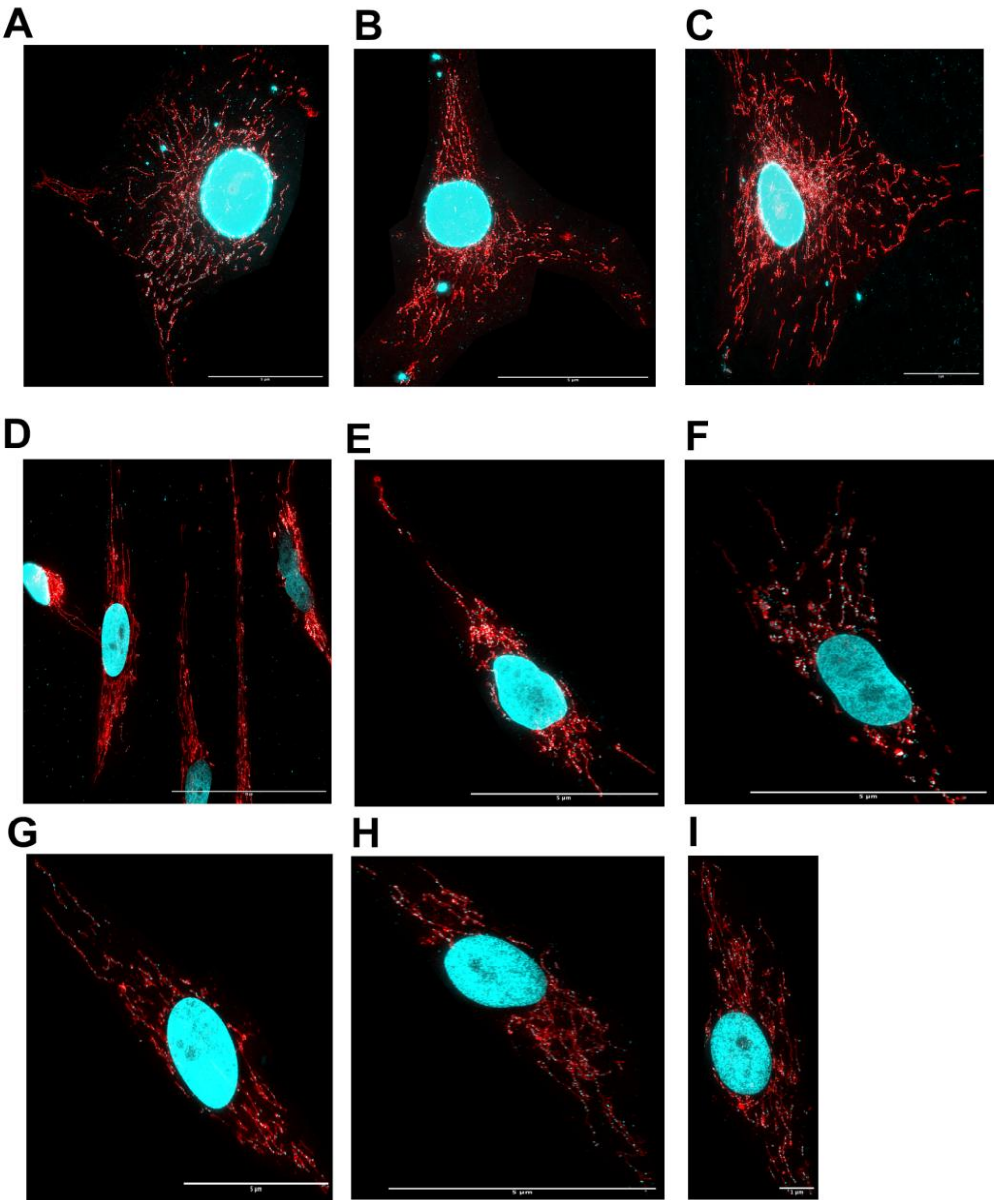


**Figure S3. Full-field microscopy images corresponding to cropped panels shown in Figure 3.** (A-C) Representative full-field microscopy images of control cells, stained with dsDNA antibody (cyan) and MitoTracker Red to visualize mtDNA nucleoids. (D-I) *MED13L* patient samples harboring P866L (D-F) and N1824M$^{fs*}$ (G-I) cell lines, stained as in (A). Full-field images are provided to demonstrate the overall staining distribution and to confirm that representative regions were chosen for presentation. Scale bars are provided for each image.

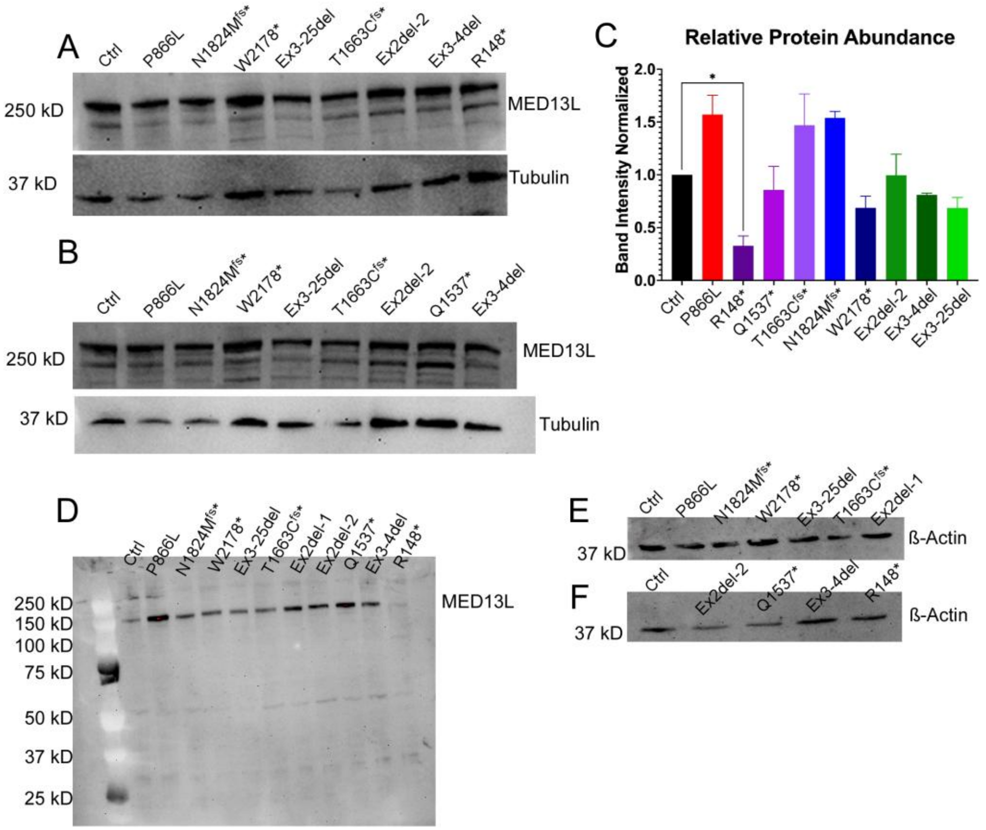


**Figure S4. Analysis of MED13L variant protein expression.** (A-B) Representative Western blot analysis of MED13L protein levels in control and *MED13L* variant fibroblast lines as indicated. β-actin levels were monitored as loading control. (C) Quantification of MED13L protein abundance from Western blot analysis. Band intensities of MED13L were normalized to loading control (β-actin) and expressed relative to control fibroblasts. (D) Western blot analysis of MED13L protein across variant fibroblast lines showing full-length and lower molecular weight species. (E-F) Loading controls corresponding to Western blot shown in (D).

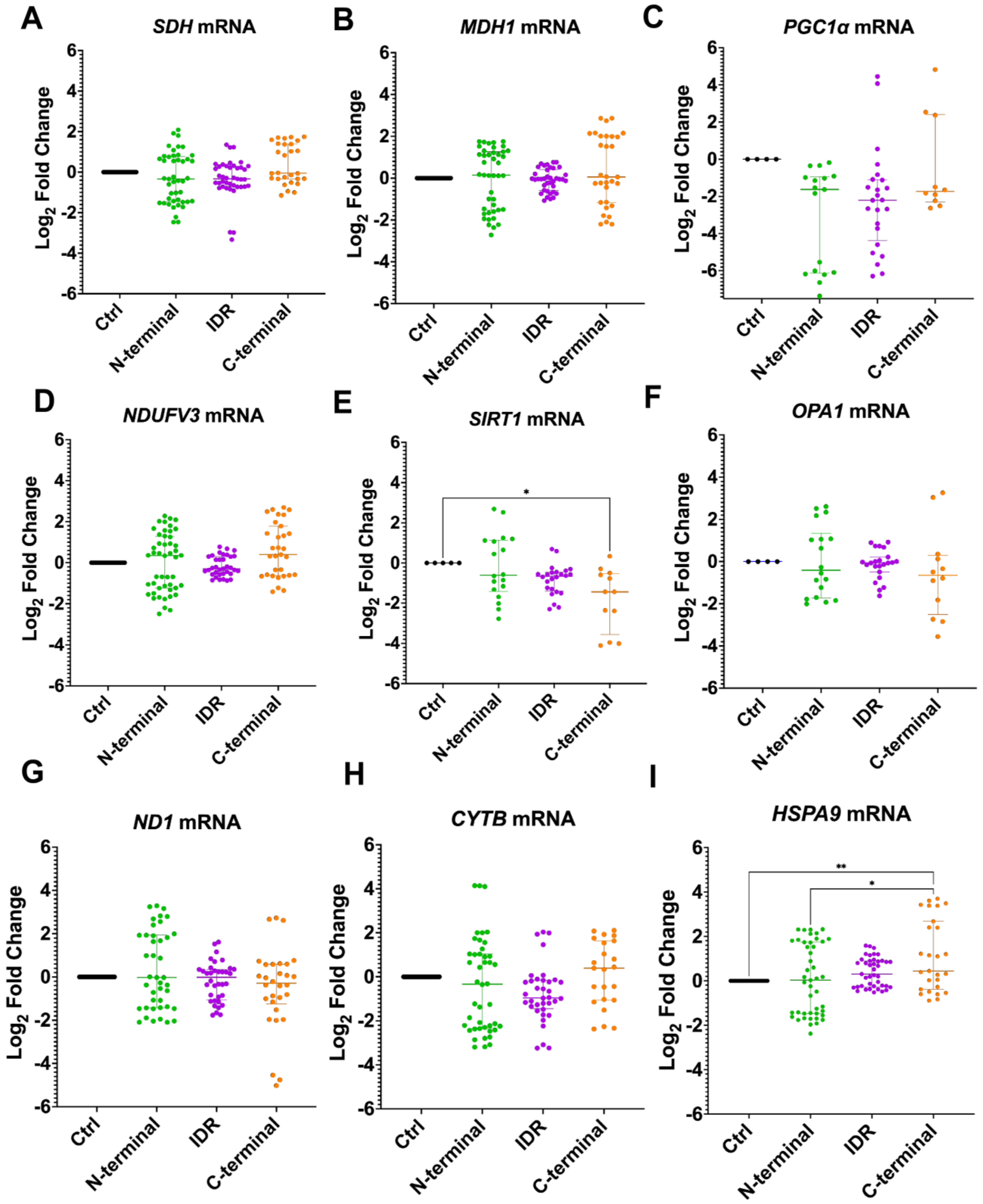
A
SDH mRNA
B
MDH1 mRNA
C
PGC1α mRNA
D
NDUFV3 mRNA
E
SIRT1 mRNA
F
OPA1 mRNA
G
ND1 mRNA
H
CYTB mRNA
I
HSPA9 mRNA
Log2 Fold Change
Ctrl
N-terminal
IDR
C-terminal
*
**

**Figure S5. Expanded analysis of mitochondrial gene expression pathways affected by *MED13L* variants.** Patient-derived fibroblast lines harboring MED13L variants were grouped according to protein domain localization and analyzed for transcriptional changes in genes associated with mitochondrial metabolism, biogenesis, dynamics, and stress responses by RT-qPCR. $Log_2$ fold change values are shown relative to control fibroblasts. (A) *SDH*, (B) *MDH1*, (C) *PGC1α*, (D) *NDUFV3*, (E) *SIRT1*, (F) *OPA1*, (G) mitochondrial-encoded *ND1*, (H) mitochondrial-encoded *CYTB*, and (I) *HSPA9* mRNA expression. ($n \geq 4$ biological replicates; technical triplicates for all genes assessed) Values represent median ± interquartile range or distributions as indicated. Statistical comparisons were performed using one-way ANOVA with Dunnett's post-hoc test. (*$p < 0.05$, **$p < 0.01$, ***$p < 0.001$, ****$p < 0.0001$).

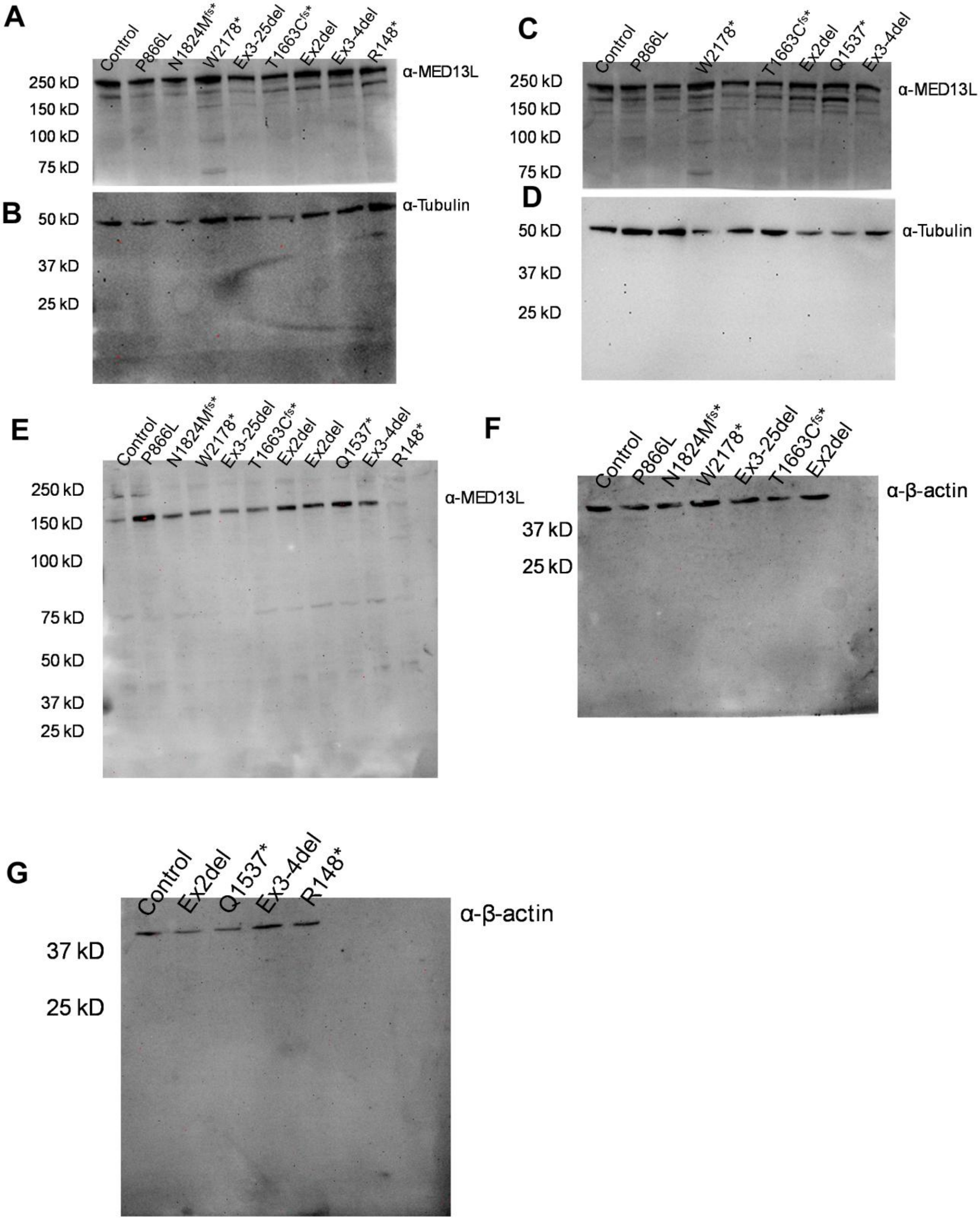

A
Control
P866L
N1824Mfs*
W2178*
Ex3-25del
T1663Cfs*
Ex2del
Ex3-4del
R148*
α-MED13L
250 kD
150 kD
100 kD
75 kD
B
α-Tubulin
50 kD
37 kD
25 kD
C
Control
P866L
W2178*
T1663Cfs*
Ex2del
Q1537*
Ex3-4del
α-MED13L
250 kD
150 kD
100 kD
75 kD
D
50 kD
37 kD
25 kD
α-Tubulin
E
Control
P866L
N1824Mfs*
W2178*
Ex3-25del
T1663Cfs*
Ex2del
Ex2del
Q1537*
Ex3-4del
R148*
250 kD
150 kD
100 kD
75 kD
50 kD
37 kD
25 kD
α-MED13L
F
Control
P866L
N1824Mfs*
W2178*
Ex3-25del
T1663Cfs*
Ex2del
α-β-actin
37 kD
25 kD
G
Control
Ex2del
Q1537*
Ex3-4del
R148*
α-β-actin
37 kD
25 kD

**Figure S6. Full membranes for Western blots found in Figure S4.** (A-B) Membrane from gradient gel (4-12%) corresponding to Figure S4A, cut in half and probed for MED13L (A) or tubulin loading control (B). (C-D) Membrane from gradient gel (4-12%) corresponding to Figure S4B, cut in half and probed for MED13L (C) or tubulin loading control (D). (E) Membrane from gradient gel (4-12%) probed for MED13L. (F) Loading control (12%) gel loaded with same samples as lanes 1-7 from gel in (E) probed for β-actin. (G) Loading control (12%) gel loaded with same samples as lanes 1, 8-12 in (E).

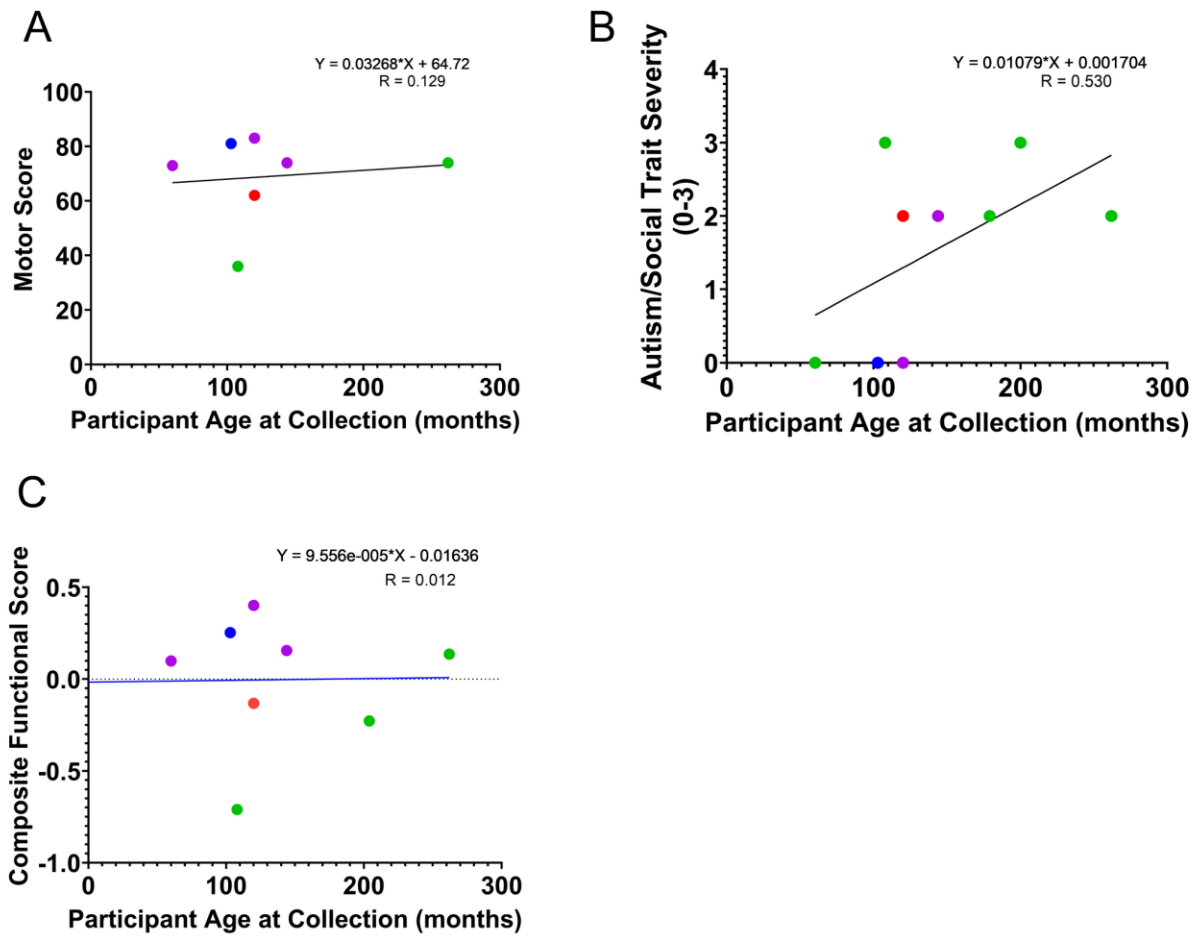


**Figure S7. Clinical outcome measures in relation to participant age.** The Vineland Motor Score (A), Autistic Severity Assessment (B) and the Composite Functional Score (C) were plotted versus participant age at the time of tissue collection. Cell lines tested for (A), (B) and (C) P866L, N1824, Ex3-25del, T1663, Q1537, L971, Ex3-4del, R148. Linear regression was performed and line of best fit shown, along with corresponding equation and R value.

**Supplemental Tables**

| Sample # | Sex | Age (approximate; months) | DNA | Protein | Mutation Type | Positional Bin | Functional Domain Bin |
|---|---|---|---|---|---|---|---|
| **1** | M | 120 | c.2597C>T | p.Pro866Leu | Missense | Early/Mid; 866 | medPIWI/IDR |
| **2** | F | 103 | c.5471delA | p.Asn1824Metfs*28 | Frameshift | Mid/Late; 1824 | C-terminal |
| **3** | M | N/A | c.6533G>A | p.Trp2178* | Nonsense | Late; 2178 | C-terminal |
| **4** | F | 204 | Del exons 3-25 | Large internal del | Multi-exon del | Early; 104 | N-terminal |
| **5** | M | 60 | c.4987_4991del | p.Thr1663Cysfs* | Frameshift | Mid; 1663 | medPIWI/IDR |
| **6** | M | 200 | Del exon 2 | Early Truncation | Exon del | Early; 43 | N-terminal |
| **7** | M | 179 | Del exon 2 | Early Truncation | Exon del | Early; 43 | N-terminal |
| **8** | M | 165 | c.4077G>A | p.Trp1359* | Nonsense | Mid; 1359 | medPIWI/IDR |
| **9** | F | 120 | c.4069C>T | p.Gln1537* | Nonsense | Mid; 1537 | medPIWI/IDR |
| **10** | M | 144 | c.2911_2914delCTGT | p.Leu971Phefs* | Frameshift | Early/Mid; 971 | N-terminal |
| **11** | F | 262 | Del exons 3-4 | Small internal del | Multi-exon del | Early; 104 | N-terminal |
| **12** | F | 108 | c.442C>T | p.Arg148* | Nonsense | Early; 148 | N-terminal |

**Table S1. Genetic and computational annotation of *MED13L* variants used in this study.** Variants were annotated using HGVS nomenclature. Position and domain bins were assigned based on canonical *MED13L* transcript structure. Multi-exon deletions removing core coding sequence were considered loss-of-function and assigned as N-terminal variants. Classifications were based on ClinVar functional domains. IDR = intrinsically disordered region.

| Gene | Application | Forward (5'-3') | Reverse (3'-5') |
|---|---|---|---|
| ***MED13L*** | qPCR | TTGTCCGACCCTACGAAAAGG | GTGCTGGGCAATCTCCACA |
| ***MED13*** | qPCR | TTGCTGGTGTCCGAATGATCT | CGTAACCGAAGACATAGCAGGAT |
| ***CCNC*** | qPCR | CCTTGCATGGAGGATAGTGAATG | AAGGAGGATACAGTAGGCAAAGA |
| ***PGC1a*** | qPCR | AGCCTCTTTGCCCAGATCTT | GGCAATCCGTCTTCATCCAC |
| ***SIRT1*** | qPCR | TCCATACCCCATGAAGTGCCTCA | GGTGTGGGTGGCAACTCTGACA |
| ***TFB1M*** | qPCR | GTGGCAGAGAGACTTGCAGCCA | GCCCACGTCCACCTCTGGTTTG |
| ***ND1*** | qPCR, mtDNA | CCCTAAAACCCGCCACATCT | GAGCGATGGTGAGAGCTAAGGT |
| ***CYTB*** | qPCR, mtDNA | CACGATTCTTTACCTTTCACTTCATC | TGATCCCGTTTCGTGCAAG |
| ***CDKN2A (p16)*** | qPCR | CCAACGCACCGAATAGTTACG | GCGCTGCCCATCATCATG |
| ***HSPA9*** | qPCR | CCCGCCACCAGGATAGCTGGAA | CTGCCACGCAGGAGTTGGTAGT |
| ***MDH1*** | qPCR | TTGTCACGACTGTGCAGCAGCG | CCCTGACGTGGTCACAGATGGC |
| ***SDHB*** | qPCR | TCGATGGGACCCAGACAAGGCT | GCATCCAATACCATGGGGCCACA |
| ***OPA1*** | qPCR | TCCAGCTGCGCAGACCATGAAT | GCTGTCCTCCGCAAAGTCATTCCA |
| ***MFN1*** | qPCR | CCGAGGAGGTGGCAAACAAAGT | GCTGGGTCTGAAGCACTAAGGCG |
| ***MFN2*** | qPCR | GCCATGTCCATGATGCCTACCCT | AAGGTGGCGCTCTCCTGGATGT |
| ***NDUFV3*** | qPCR | TGCTGTGGCCCTGCTTGGTG | ACACCTGGGCTTCCTGGAGCAT |
| ***PK*** | Reference | AGCCCAAATGGCCTTGAA | AGAGACAGAATGCCAGTGAGC |
| ***GAPDH*** | Reference | GGTTGTCTCCTGCGACTTCA | CCCTAGGCCCCTCCTGTTAT |

**Table S2. Oligonucleotide sequences used for qPCR and mtDNA copy number quantification.** Forward and reverse primer sequences used for RT-qPCR analysis of nuclearencoded and mitochondrial-encoded transcripts, as well as mtDNA copy number quantification. Reference genes used for normalization are indicated.

| **Construct** | **Simons Searchlight Variable** | **RARE-X Variable** | **Harmonized Variable** | **Tiering/Binning** |
|---|---|---|---|---|
| Adaptive Function | ABC Standard Score | vi3_adaptive_composite | Adaptive Severity | >85 mild; 70-84 borderline; <70 moderate/severe |
| Communication | Vineland communication | vi3_comm_domain | Communication Severity | Same as above |
| Motor | Motor Domain | vi3_motor_domain | Motor Severity | Same as above |
| SCQ | SCQ total | | SCQ Tier | <10=0; 10-14=1; 15-21=2; >21=3 |
| SRS | srs_total_t | | SRS Tier | <60=0; 60-65=1; 66-75=2; >75=3 |
| Socialization | | vi3_socialization | Social Severity | >85=0; 70-84=1; 55-69=2; <55=3 |

**Table S3. Harmonized Behavioral Variable Definitions and Scoring**

Variables from Vineland-3, Social Communication Questionnaire (SCQ), Social Responsiveness Scale (SRS), and caregiver- or clinician-reported autism spectrum disorder (ASD) diagnoses were harmonized across registries to generative adaptive function, communication, motor, social impairment, and autism-associated behavioral severity metrics used for downstream genotype-phenotype analyses. Tier classifications and score cutoffs used for severity assignments are shown. Composite Functional Scores were generated from normalized adaptive behavior, communication, and motor domain scores, with inversion of Adaptive Behavior Composite (ABC) z-score prior to averaging to maintain consistent impairment directionality across variables.

| **Individual** | **ABC Score** | **ABC Score Normalized & Inverted** | **Communication Raw** | **Communication Normalized** | **Motor Score Raw** | **Motor Score Normalized** | **Composite Score** |
|---|---|---|---|---|---|---|---|
| **#1** | **62** | **0.386** | **59** | **-0.345** | **62** | **-0.436** | **-0.132** |
| **#2** | **73** | **-0.086** | **70** | **0.098** | **81** | **0.748** | **0.253** |
| **#3** | **n/a** | **n/a** | **n/a** | **n/a** | **n/a** | **n/a** | **n/a** |
| **#4** | **114** | **-1.844** | **102** | **1.388** | **n/a** | **n/a** | **-0.228** |
| **#5** | **75** | **-0.171** | **73** | **0.219** | **73** | **0.249** | **0.099** |
| **#6** | **n/a** | **n/a** | **n/a** | **n/a** | **n/a** | **n/a** | **n/a** |
| **#7** | **n/a** | **n/a** | **n/a** | **n/a** | **n/a** | **n/a** | **n/a** |
| **#8** | **n/a** | **n/a** | **n/a** | **n/a** | **n/a** | **n/a** | **n/a** |
| **#9** | **74** | **-0.129** | **79** | **0.461** | **83** | **0.873** | **0.402** |
| **#10** | **71** | **0** | **n/a** | **n/a** | **74** | **0.311** | **0.156** |
| **#11** | **71** | **0** | **70** | **0.098** | **74** | **0.311** | **0.137** |
| **#12** | **28** | **1.844** | **20** | **-1.917** | **36** | **-2.057** | **-0.710** |

**Table S4. Individual Participant Behavioral and Functional Scores.** Individual participant adaptive behavior, communication, and motor scores used to generate Composite Functional Scores. Raw values, normalized values, and inverted adaptive behavior scores are shown. Composite scores were calculated from normalized measures to facilitate cross-domain comparisons of functional impairment.

| Individual | Autism/Social Trait Tier | SRS | SCQ | Vineland Socialization |
|---|---|---|---|---|
| #1 | 2 | n/a | 18 | n/a |
| #2 | 0 | n/a | 5 | n/a |
| #3 | n/a | n/a | n/a | n/a |
| #4 | 0 | n/a | 8 | n/a |
| #5 | 0 | n/a | n/a | 85 |
| #6 | 3 | 84 | 19 | n/a |
| #7 | 2 | 73 | 14 | n/a |
| #8 | n/a | n/a | n/a | n/a |
| #9 | 0 | 59 | 13 | n/a |
| #10 | 2 | 68 | 5 | 74 |
| #11 | 2 | n/a | 18 | n/a |
| #12 | 3 | n/a | 32 | n/a |

**Table S5. Autism/Social Trait Raw Scores used for Autism/Social Trait Severity Tiering.** Individual participant scores used for autism/social trait severity classification, including Social Responsiveness Scale (SRS), Social Communication Questionnaire (SCQ), Vineland Socialization scores, and derived Autism/Social Trait severity tier assignments. Missing values indicate data unavailable for a given participant.
